# Osmotic stress-induced, rapid clustering of IGF2BP proteins nucleates stress granule assembly

**DOI:** 10.1101/846659

**Authors:** Wei-jie Zeng, Chuxin Lu, Yuanyuan Shi, Xinxin Chen, Jie Yao

**Author notes:** Present Address: Allen Institute for Cell Science, Seattle, WA, 98109, USA.

## Abstract

Stress granules (SGs) are formed in the cytoplasm by liquid-liquid phase separation (LLPS) of translationally-stalled mRNA and RNA-binding proteins during stress response. Understanding the mechanisms governing SG assembly requires imaging SG formation in real time. Here we used live cell imaging and super-resolution imaging to visualize SG assembly in human cells. We found that IGF2BP proteins formed microscopically visible clusters almost instantaneously upon osmotic stress, prior to the recruitment of G3BP1 and TIA1. The rapid clustering of IGF2BP1 was ATP-independent and was mediated by its KH3/4 di-domains and an intrinsically disordered region (IDR), whereas ATP depletion inhibited the recruitment of G3BP1 and TIA1. Moreover, we detected cytoplasmic clusters of IGF2BP1 below the optical resolution in normal cells and found IGF2BP1 forming a dense granule associated with multiple clusters of poly(A) mRNA in mature SGs. Thus, ATP-independent, rapid clustering of IGF2BP nucleates SG assembly during osmotic stress.

## Introduction

In eukaryotic cells, membraneless organelles perform important roles in normal cell functions and during response to external stimuli (Banani, Lee, Hyman, & Rosen, 2017; Kedersha & Anderson, 2007; Shin & Brangwynne, 2017). For instance, the nucleolus is the location of ribosomal RNA transcription and processing, and SGs are formed upon various environmental stress conditions as the storage place for translationally-stalled mRNAs (Panas, Ivanov, & Anderson, 2016; Protter & Parker, 2016). Membraneless organelles are formed through LLPS, in which macromolecules including RNA and proteins coalesce and form microscopically-visible clusters in vitro or in vivo (Banani et al., 2017; Shin & Brangwynne, 2017). Multiple types of macromolecular interactions were proposed to mediate LLPS in the cell, including multivalent interactions among structured protein domains or among IDRs, RNA-protein interactions and RNA-RNA interactions (Banani et al., 2017; Panas et al., 2016; Van Treeck & Parker, 2018).

SGs are formed during various stress conditions, such as oxidative stress, osmotic stress, heat shock, etc. Major components of SGs include untranslated mRNA, ribosomal protein subunits and RNA-binding proteins (Aulas et al., 2017; Buchan & Parker, 2009; Jain et al., 2016; Khong et al., 2017). Proteomic analyses and subsequent studies have identified additional protein factors associated with SGs such as ATPases, kinases and RNA methyltransferases, as well as key proteins regulating SG assembly and disassembly (Jain et al., 2016; Mahboubi, Koromilas, & Stochaj, 2016; Markmiller et al., 2018; Ries et al., 2019; Turakhiya et al., 2018; Wippich et al., 2013). Mutations of many SG proteins such as VCP/p97 was found in neurodegenerative diseases including amyotrophic lateral sclerosis (Buchan, Kolaitis, Taylor, & Parker, 2013; Turakhiya et al., 2018). Visualizing how SGs are assembled and organized in a living cell is essential for understanding SG regulatory pathways and LLPS processes in general.

Light microscopy is an ideal approach to reveal both the real-time dynamics of macromolecules undergoing LLPS and the molecular organization of membraneless organelles (Van Treeck & Parker, 2019). Wheeler et al used live cell imaging to probe the dynamics of SG assembly and disassembly processes under oxidative stress (Wheeler, Matheny, Jain, Abrisch, & Parker, 2016). Single molecule RNA fluorescence in situ hybridization (FISH) has revealed the assembly kinetics of different endogenous mRNA and their orientations within SGs (Khong & Parker, 2018). Live cell imaging of single RNA molecules has revealed the kinetics of RNA recruitment to single SGs (Moon et al., 2019; Wilbertz et al., 2019). The general concept arising from previous studies was that SG assembly proceeds as a core-shell model: translationally-stalled mRNA serves as a core component for SG assembly and subsequently recruits additional SG proteins (Protter & Parker, 2016). Consistent with the core-shell model, SGs were observed to first disintegrate into smaller particles upon stress removal and then become completely disassembled (Wheeler et al., 2016). Although oxidative stress has been widely used for imaging SG assembly, SG assembly dynamics and SG organizations in other stress conditions have largely escaped analysis.

In this study, we examined SG assembly kinetics in human cells by live cell imaging and super-resolution microscopy, and compared SG assembly kinetics under osmotic stress versus oxidative stress conditions. Remarkably, we observed distinct recruitment kinetics of SG proteins during osmotic stress. IGF2BP proteins formed microscopically-visible clusters in the cytoplasm almost instantaneously after osmotic shock, while several other SG proteins such as G3BP1/TIA1 were recruited to IGF2BP1-marked granules at a later stage. Both the KH3/KH4 di-domain of IGF2BP1 and an IDR between its RRM domain and KH1 domain contributed to its rapid clustering upon osmotic stress. Furthermore, super-resolution microscopy revealed that IGF2BP1 existed as small clusters (100-200 nm in diameter) in the cytoplasm prior to stress, rapidly coalesced into larger clusters (250-500 nm in diameter) upon osmotic stress, and formed a “core-like” granule within mature SGs during extended stress. Taken together, our studies uncovered a novel SG assembly pathway in which the rapid clustering of IGF2BP proteins nucleates SG assembly during osmotic stress.

## Results

### Rapid clustering of IGF2BP1 preceded the recruitment of G3BP1 and TIA1 to SGs during osmotic stress

IGF2BP1 (also named as IMP1 or ZBP1) is a key RNA-binding protein and an SG component (Bell et al., 2013). Previous studies have reported that IGF2BP1 was recruited along with other proteins to SGs during oxidative stress (Bley et al., 2015; Markmiller et al., 2018; Niewidok et al., 2018; Stohr et al., 2006). During oxidative stress, multiple SG proteins were detected to coalesce into SGs at about 15 min after cell treatment with sodium arsenite (Moon et al., 2019; Wheeler et al., 2016). Although mRNA of longer lengths was found to localize to SGs after a delay time upon stress (Khong & Parker, 2018), it remains unknown whether different SG proteins were recruited to SGs as a single step or as multiple steps. In this work, we first examined the recruitment kinetics of three SG proteins, IGF2BP1, G3BP1 and TIA1 during osmotic stress or oxidative stress in living human U2OS cells. Surprisingly, we found that GFP-IGF2BP1 or mCherry-IGF2BP1 formed microscopically-visible clusters in the cytoplasm very rapidly after applying osmotic stress (Figure 1A, Figure 1 – figure supplement 1A, Video 1). Because we had to pause image acquisition for approximately 40 seconds to allow for the application of culture media containing 375 mM sorbitol to cells, we estimated that the time delay between applying osmotic stress and IGF2BP1 forming clusters was less than 40 seconds. Although IGF2BP1 was predominantly localized in the cytoplasm, some IGF2BP1 clusters were also formed in the nucleus (Figure 1A). Furthermore, we observed rapid clustering of IGF2BP1 upon applying hypertonic culture media to cells (Figure 1 – figure supplement 1B, Video 2), indicating that the rapid clustering of IGF2BP1 was not induced only by sorbitol but was instead a cellular response to increased osmolarity of the culture environment.

**Figure 1.**
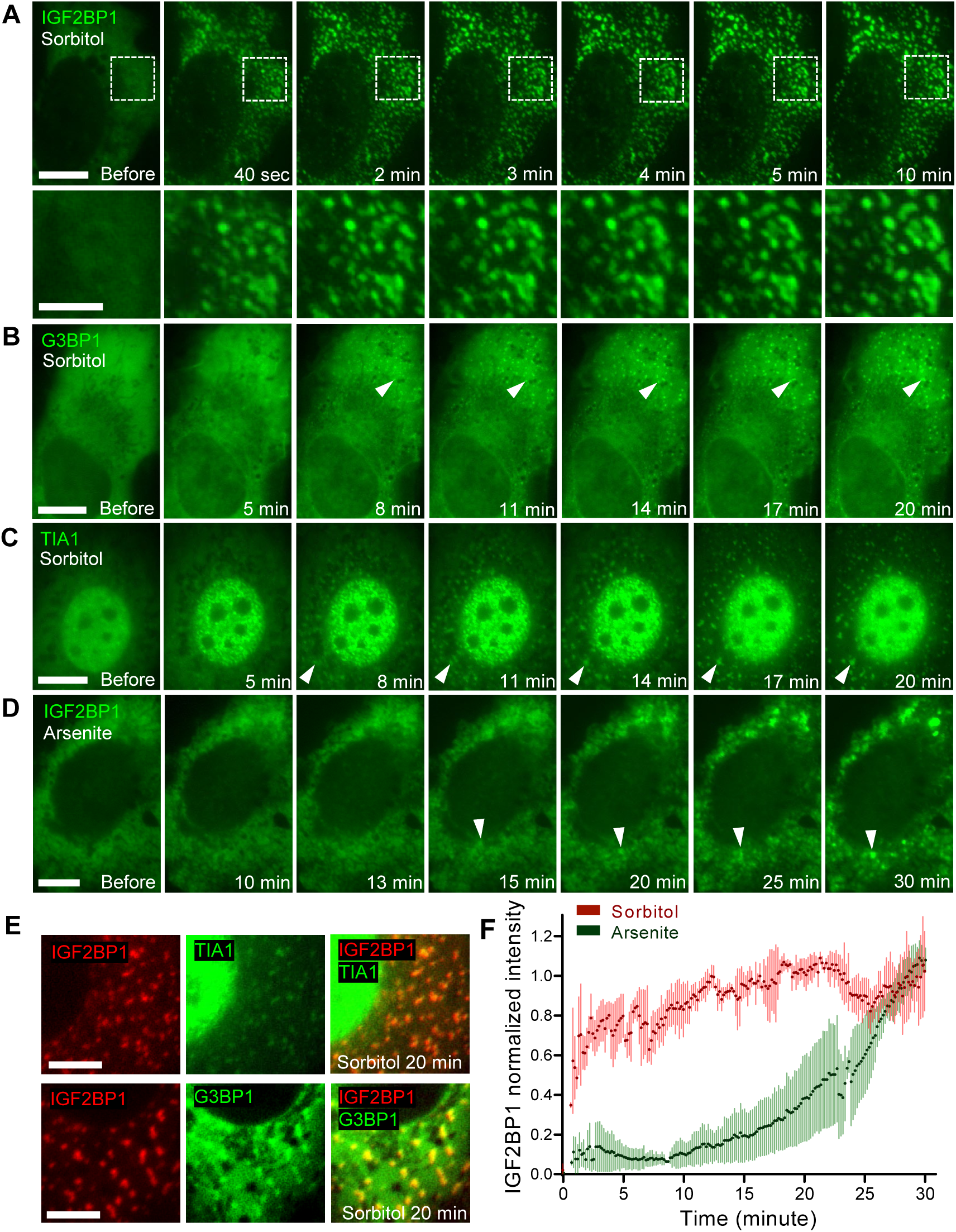
Rapid clustering of IGF2BP1 upon osmotic stress precedes the recruitment of other SG proteins. **(A)** Time-lapse images of GFP-IGF2BP1 in U2OS cells treated with 375 mM sorbitol. Scale bars: 10 μm (upper row) and 5 μm (lower row). The lower row displays the enlarged images of areas marked by dashed white lines in the upper row. **(B-D)** Time-lapse images of GFP-tagged G3BP1(B), TIA1 (C) or IGF2BP1(D) in U2OS cells treated with 375mM sorbitol or with 0.5mM sodium arsenite, as indicated in the figures. Scale bars are 10 μm. Arrowheads indicate SGs during assembly. Time intervals between applying stressors and image acquisition are noted on each image. **(E)** Dual-color fluorescence images of U2OS cells co-expressing mCherry-IGF2BP1 and TIA1-GFP (upper) or GFP-G3BP1 (lower) after treatment with 375mM sorbitol for 20 min. Scale bars are 5 μm. **(F)** Normalized fluorescence intensities of GFP-IGF2BP1 at SG regions during treatment with 375mM sorbitol (n = 4) or 0.5mM sodium arsenite (n = 3). Interval between adjacent time points: 10 seconds. Whiskers indicate standard deviations at each time point.

The small clusters marked by GFP-IGF2BP1 grew in size and coalesced into larger granules within 5 min after sorbitol treatment (Figure 1A). In contrast, GFP-G3BP1 or TIA1-GFP didn’t exhibit such a rapid clustering during osmotic stress (Figure 1B-C, Video 3, 4). We found that GFP-G3BP1 and TIA1-GFP were recruited to SGs at approximately 8 min following sorbitol treatment (Figure 1 – figure supplement 2C). For comparison, we examined the recruitment of these three SG proteins under oxidative stress. IGF2BP1 was assembled into SGs at 15-20 min after sodium arsenite treatment (Figure 1D, Video 5), about the same time when G3BP1 and TIA1 were assembled into SGs (Figure 1-figure supplement 2A-C, Video 6, 7). Importantly, IGF2BP1 was colocalized with TIA1 and G3BP1 in SGs during both osmotic stress (Figure 1E) and oxidative stress (Figure 1 – figure supplement 2D), indicating that IGF2BP1 is an integral SG component during both stress conditions.

Next, we quantified fluorescence intensities of GFP-IGF2BP1 at SG regions during the time course of osmotic stress and oxidative stress. GFP-IGF2BP1 fluorescence intensity at SGs (Figure 1F) was dramatically increased at 40 sec after sorbitol treatment and reached ∼80 % of maximum intensity at approximately 3 min after stress. In contrast, GFP-IGF2BP1 fluorescence intensity at SGs started to substantially increase at ∼15 min after treatment with 0.5 mM sodium arsenite and reached the highest level at the end of the time course (about 30 min after stress, Figure 1F). Taken together, our studies indicated that during oxidative stress, different SG proteins were assembled into SGs at a rather uniform rate and a delay time (usually 15 min) was likely required prior to SG assembly, while during osmotic stress, IGF2BP1 formed clusters almost instantaneously upon stress and other SG proteins were recruited to SGs at a later stage (about 8 min after stress). Our findings thus revealed a novel multiphase SG assembly kinetics during osmotic stress that was initiated by the rapid clustering of IGF2BP1.

### Rapid clustering of IGF2BP1 was ATP-independent

Previous studies showed that ATP depletion blocked SG formation during oxidative stress (Jain et al., 2016). To examine the ATP-dependence of SG assembly during osmotic stress, we treated U2OS cells with culture media containing sorbitol, 2-deoxyglucose (2DG) and carbonyl cyanide m-chlorophenyl hydrazone (CCCP). We found that GFP-IGF2BP1 formed visible clusters at 40 seconds after treated with media containing 375 mM sorbitol and ATP depletion reagents, but the clusters stopped growing in size as stress continued (Figure 2A, Video 8). Instead, GFP-G3BP1 formed weak clusters in the cytoplasm at ∼20 min after osmotic stress and ATP depletion (Figure 2B, Video 9), much more slowly than during osmotic stress alone (∼8 min, see Figure 1B). The clustering of TIA1-GFP was barely visible in the cytoplasm but was more pronounced in the nucleus (Figure 2C, Video 10). Therefore, the rapid clustering of IGF2BP1 in the cytoplasm (and TIA1-GFP clustering in the nucleus) upon osmotic stress was ATP-independent and likely resulted from diffusion-driven macromolecular interactions, whereas subsequent growth of SGs accompanied by the recruitment of G3BP1 and TIA1 was an ATP-dependent, energy-driven process.

**Figure 2.**
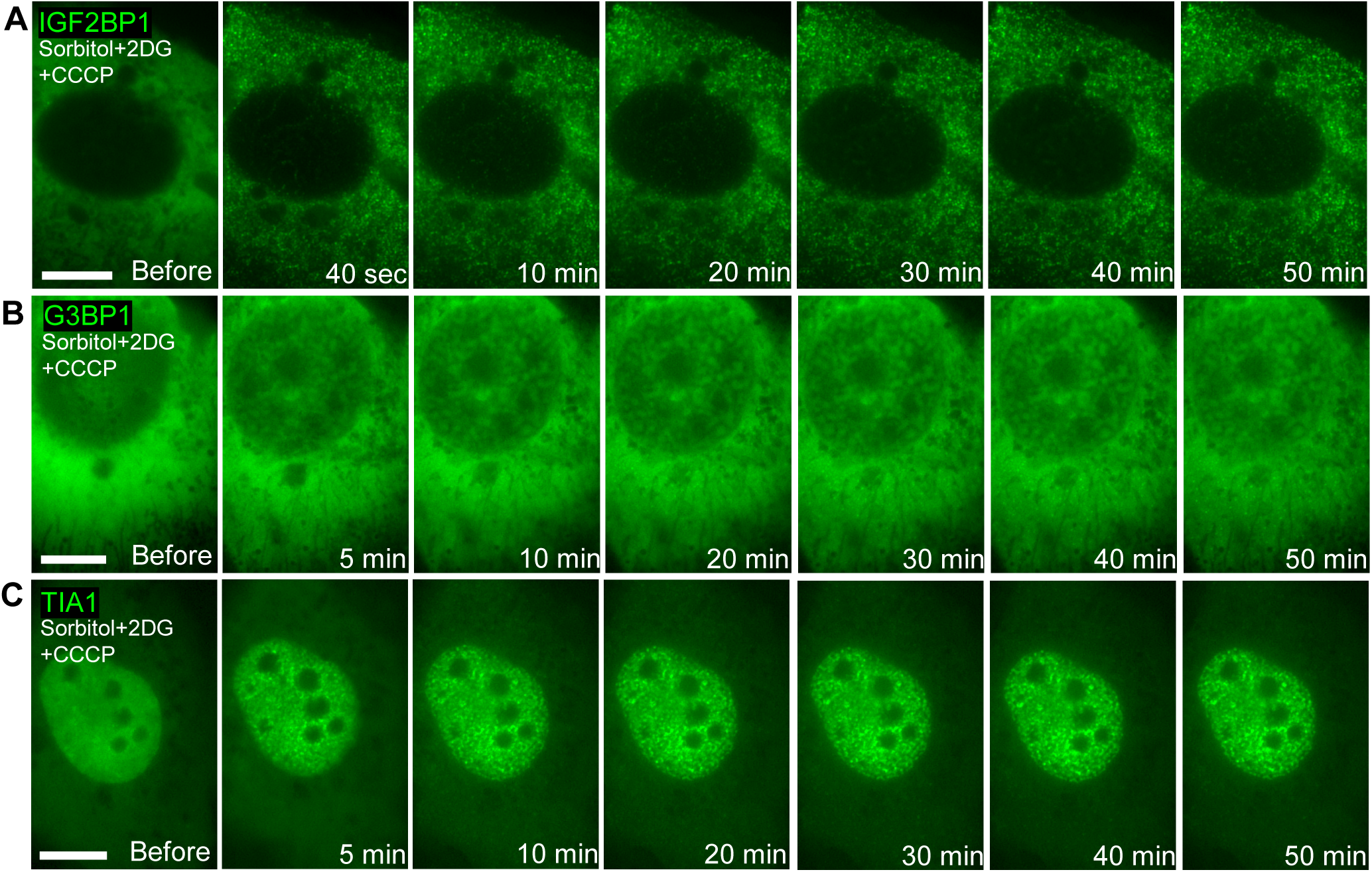
Clustering of different SG proteins during osmotic stress and ATP depletion. **(A-C)** Time-lapse fluorescence images of GFP-IGF2BP1 (A), GFP-G3BP1 (B) and TIA1-GFP (C) in U2OS cells treated with 375mM sorbitol, 200mM 2-deoxyglucose and 100µM CCCP. Time intervals between applying osmotic stress and image acquisition are noted on each image. All scale bars are 10 μm.

### Members of the IGF2BP protein family were all rapidly clustered after osmotic stress

The human IGF2BP protein family includes three homologous genes, IGF2BP1, IGF2BP2 and IGF2BP3, all of which encode proteins with highly similar amino acid sequences (Bell et al., 2013). Next, we examined the recruitment kinetics of IGF2BP2 and IGF2BP3 to SGs during osmotic stress. Similar to IGF2BP1, both GFP-IGF2BP2 and mCherry-IGF2BP3 showed significant clustering in the cytoplasm at 40 seconds after osmotic stress and formed larger SGs during extended osmotic stress (∼10 min, Figure 3A). We confirmed that the clustering of IGF2BP2 and IGF2BP3 occurred simultaneously as IGF2BP1 in the same cell at 40 seconds after osmotic stress (Figure 3B, C). Furthermore, we found that IGF2BP1, IGF2BP2 and IGF2BP3 were colocalized in the SGs (Figure 3A-C). These results demonstrated that all three members of the IGF2BP protein family were rapidly clustered in the cytoplasm after osmotic stress to initiate SG formation in human cells.

**Figure 3.**
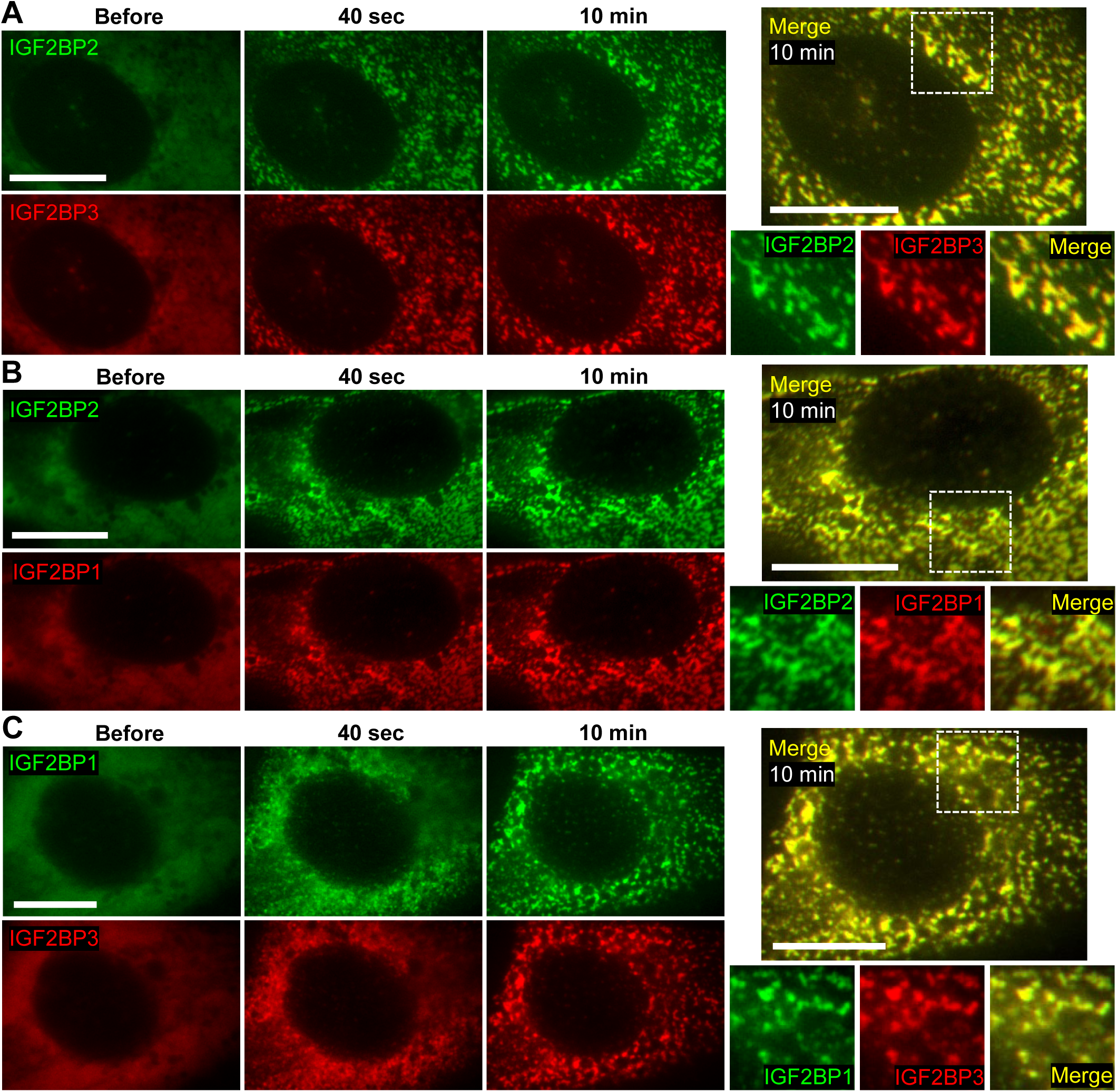
Members of the IGF2BP protein family were all rapidly clustered after osmotic stress and were colocalized. **(A-C)** Time-lapse dual-color fluorescence images of GFP-IGF2BP2/mCherry-IGF2BP3 (A), GFP-IGF2BP2/mCherry-IGF2BP1 (B), and GFP-IGF2BP1/mCherry-IGF2BP3 (C) in U2OS cells treated with 375 mM sorbitol. Time intervals between applying osmotic stress and image acquisition are noted in each image. Scale bars are 10 μm.

### KH3/4 di-domain and an IDR mediate the rapid clustering of IGF2BP1 after osmotic stress

Next, we analyzed the protein domains mediating the rapid clustering of IGF2BP1 upon osmotic stress. Previous studies found that in *Xenopus* (Git & Standart, 2002) and chicken (Farina, Huttelmaier, Musunuru, Darnell, & Singer, 2003), the K homology (KH) domain of IGF2BP1 was necessary for mediating protein oligomerization in vitro and granule formation in cells. The human IGF2BP1 protein contains four KH domains as RNA-binding modules (Chao et al., 2010; Nielsen, Nielsen, Kristensen, Koch, & Christiansen, 2002). Whether these domains regulate the rapid clustering of IGF2BP1 is not known. Therefore, we generated DNA constructs expressing different human IGF2BP1 truncation fragments according to the described domains (Chao et al., 2010) (Figure 4A), transfected these constructs into U2OS cells and performed live cell imaging during osmotic stress. Care was taken during imaging to maintain identical illumination conditions and to ensure imaging cells with similar expression levels of GFP-fusion proteins.

**Figure 4.**
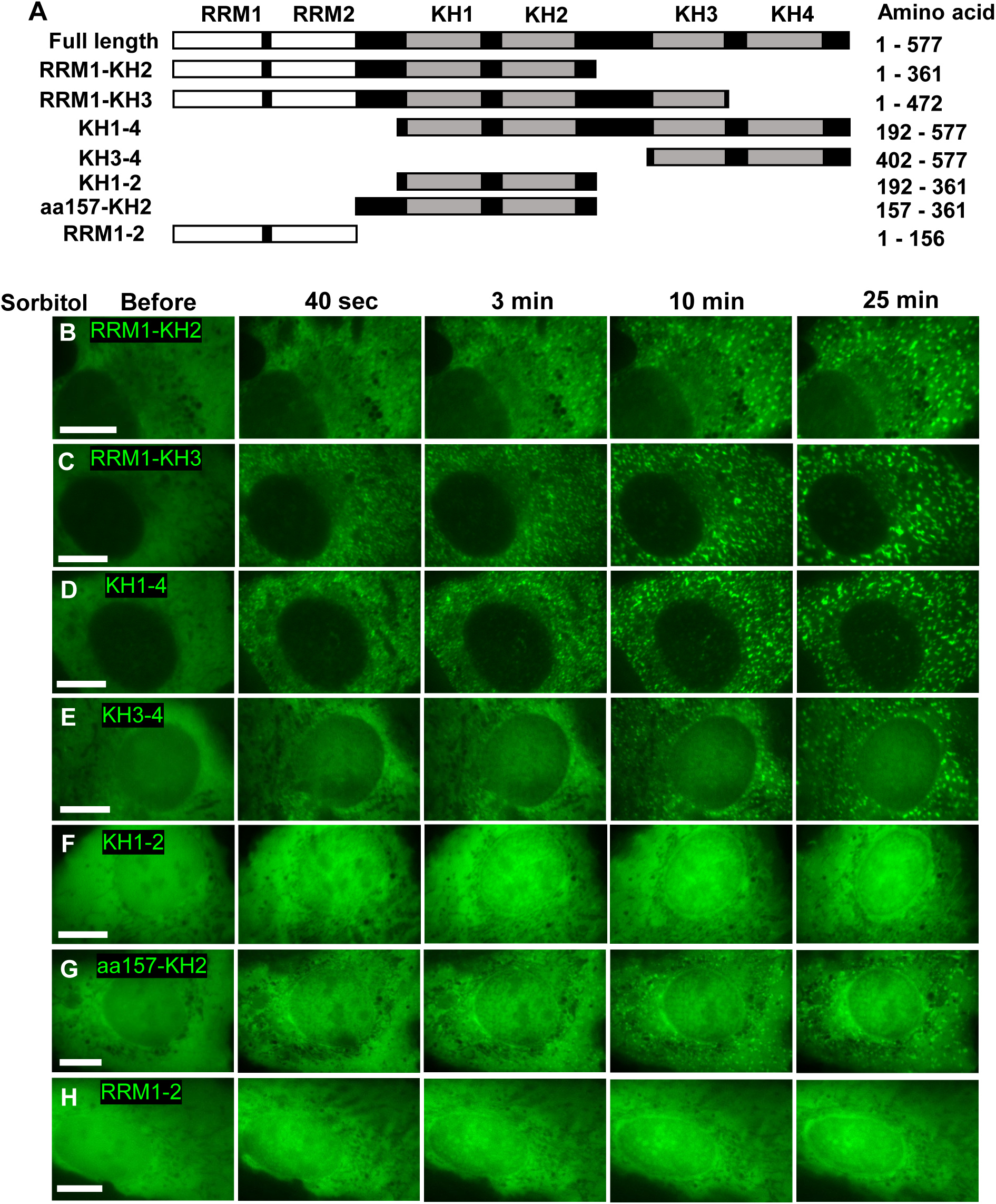
Identifying protein domains mediating the rapid clustering of IGF2BP1 during osmotic stress. **(A)** Schematic representation of IGF2BP1 and its truncation mutants examined in this study. Black boxes represent linker regions between the protein domains. The amino acid residues of each mutant are shown to the right of each diagram. The diagrams were not drawn to scale. **(B-H)** Time-lapse images of GFP-tagged IGF2BP1 truncation mutants (described in A) in U2OS cells treated with 375mM sorbitol. Time intervals between applying osmotic stress and image acquisition are noted on each image. All scale bars are 10 μm.

First, we observed a rapid clustering (within 40 seconds) of RRM1-KH2, RRM1-KH3, KH1-4 and KH3-4 fragments in U2OS cells upon osmotic stress (Figure 4B-E, Video 11-14). Clusters formed by these fragments coalesced into larger granules as stress proceeded. Next, we found that the KH1-2 fragment (Figure 4F, Video 15) and the RRM1-2 fragment (Figure 4H, Video 17) didn’t rapidly form clusters upon osmotic stress, while several small aggregates of KH1-2 fragments were formed in the cytoplasm at the later stage (25 min or later after stress). Therefore, the KH3/4 di-domain of IGF2BP1 was sufficient to induce rapid clustering but was not required for cluster formation upon osmotic stress. Either the RRM1/RRM2 domain or the KH1/2 domain did not promote rapid clustering during osmotic stress. However, because the RRM1-KH2 fragment rapidly formed clusters upon osmotic stress, we hypothesized that an amino acid region located within this fragment played key roles in promoting cluster formation.

Interactions among IDRs were proposed as a major mechanism underlying LLPS in cells (Banani et al., 2017; Molliex et al., 2015; Protter & Parker, 2016; Shin & Brangwynne, 2017). Interestingly, when we examined an IGF2BP1 fragment (aa157-KH2) containing the KH1/2 domain and an upstream linker region containing an IDR (amino acid sequence: EQIAQGPENGRRGGFGSRGQPRQGSPVAAGAPAKQ), we found that this fragment promptly formed clusters (within 3 min) after sorbitol treatment (Figure 4G, Video 16). Even though the aa157-KH2 fragment was not as rapidly clustered (∼3 min) as other IGF2BP1 fragments examined (within 40 sec), this short IDR region substantially accelerated the clustering of KH1/2 di-domain (3 min in aa157-KH2 vs. 25 min in KH1-2) during osmotic stress. Additionally, the KH1/2 di-domain and/or the region between KH1/2 and KH3/4 di-domains might enhance the clustering (compare Figure 4D and 4E). In conclusion, we identified that multiple amino acid regions of human IGF2BP1, in particular the KH3/4 di-domain and an IDR region upstream of the KH1/2 di-domain, played major roles in its rapid clustering upon osmotic stress.

### Distinct incorporation of IGF2BP1 fragments into SGs during osmotic stress and oxidative stress

To further examine the roles of IGF2BP1 protein domains in SG assembly, we cotransfected IGF2BP1 truncation mutants (described above) and mCherry-G3BP1 into U2OS cells and examined the incorporation of these mutants into G3BP1-labeled SGs during osmotic stress and oxidative stress. At 30 min after osmotic stress, all mutants except RRM1-2 were colocalized with SGs marked by mCherry-G3BP1 (Figure 5A). Therefore, self-associations mediated by the KH3/4 di-domain and its upstream regions might help to guide subsequent recruitment of other SG proteins such as G3BP1.

**Figure 5.**
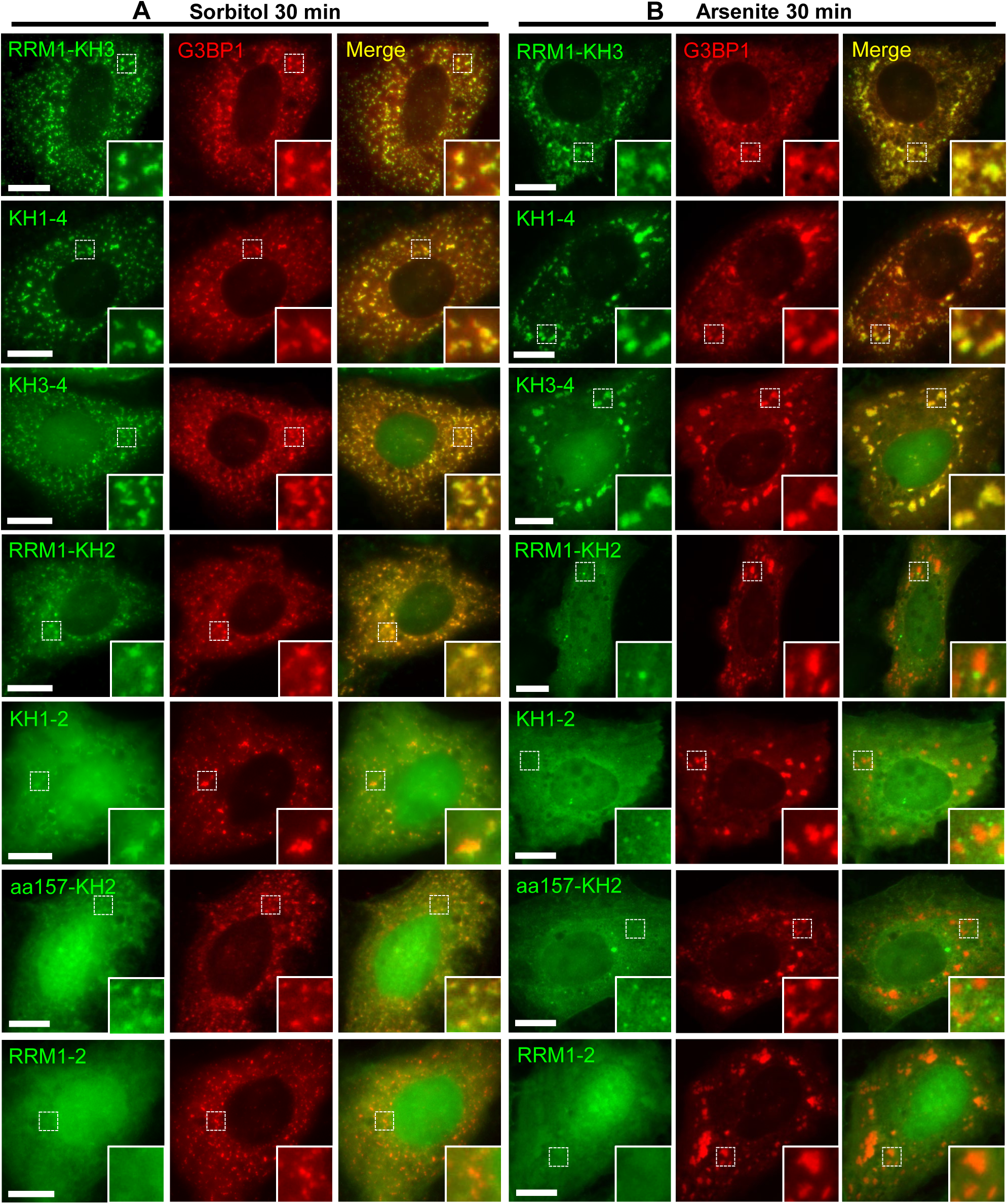
Differential colocalizations of IGF2BP1 truncation mutants with SG marker protein G3BP1 during osmotic stress or oxidative stress. (A, B) Dual-color fluorescence images of U2OS cells co-expressing mCherry-G3BP1 and GFP-tagged IGF2BP1 truncation mutants (described in Figure 4A) after treatment with 375mM sorbitol (A) or 0.5mM sodium arsenite (B) for 30 min. The inset at the lower-right corner of each image is the enlarged image of the area bordered by the dashed line. All scale bars are 10 μm.

During oxidative stress, we found that all GFP-tagged IGF2BP1 fragments except RRM1-2 formed cytoplasmic punta approximately 15 min after arsenite treatment (Figure 4 – figure supplement 1). At 30 min after oxidative stress, clusters formed by RRM-KH3, KH1-4 and KH3-4 fragments were colocalized with mCherry-G3BP1 (Figure 5B). Interestingly, clusters formed by IGF2BP1 fragments lacking the KH3/4 di-domain (such as RRM-KH2, KH1-2 and aa157-KH2 fragments) were not colocalized with SGs marked by mCherry-G3BP1 (Figure 5B). These results suggested that the KH3/4 di-domain, but not its upstream regions, mediated the incorporation of IGF2BP1 into SGs during oxidative stress. It is possible that the RNA-binding activity of KH3/4 di-domains drives the recruitment of IGF2BP1 into SGs during oxidative stress.

### Poly(A) mRNA was rapidly associated with IGF2BP1 clusters during osmotic stress

One major function of SGs is to serve as a storage site for translationally-stalled mRNAs (Buchan & Parker, 2009; Kedersha & Anderson, 2007; Panas et al., 2016; Protter & Parker, 2016). Therefore, we were interested in understanding the assembly process of mRNA molecules into SGs during osmotic stress. First, we generated a stable U2OS cell line expressing GFP-IGF2BP1 in order to detect IGF2BP1 and poly(A) mRNA in the same cell. We confirmed the rapid clustering of GFP-IGF2BP1 upon osmotic stress in this stable cell line (Figure 6 – figure supplement 1A). About 20-30% of the cells in this cell line expressed GFP-IGF2BP1 (Figure 6 – figure supplement 1B). Western blot and titration analysis showed that GFP-IGF2BP1 was expressed at a level at about 10% of the endogenous IGF2BP proteins (Figure 6 – figure supplement 1C, data not shown). Thus, the level of overexpressed GFP-IGF2BP1 was lower than that of endogenous IGF2BP proteins.

**Figure 6.**
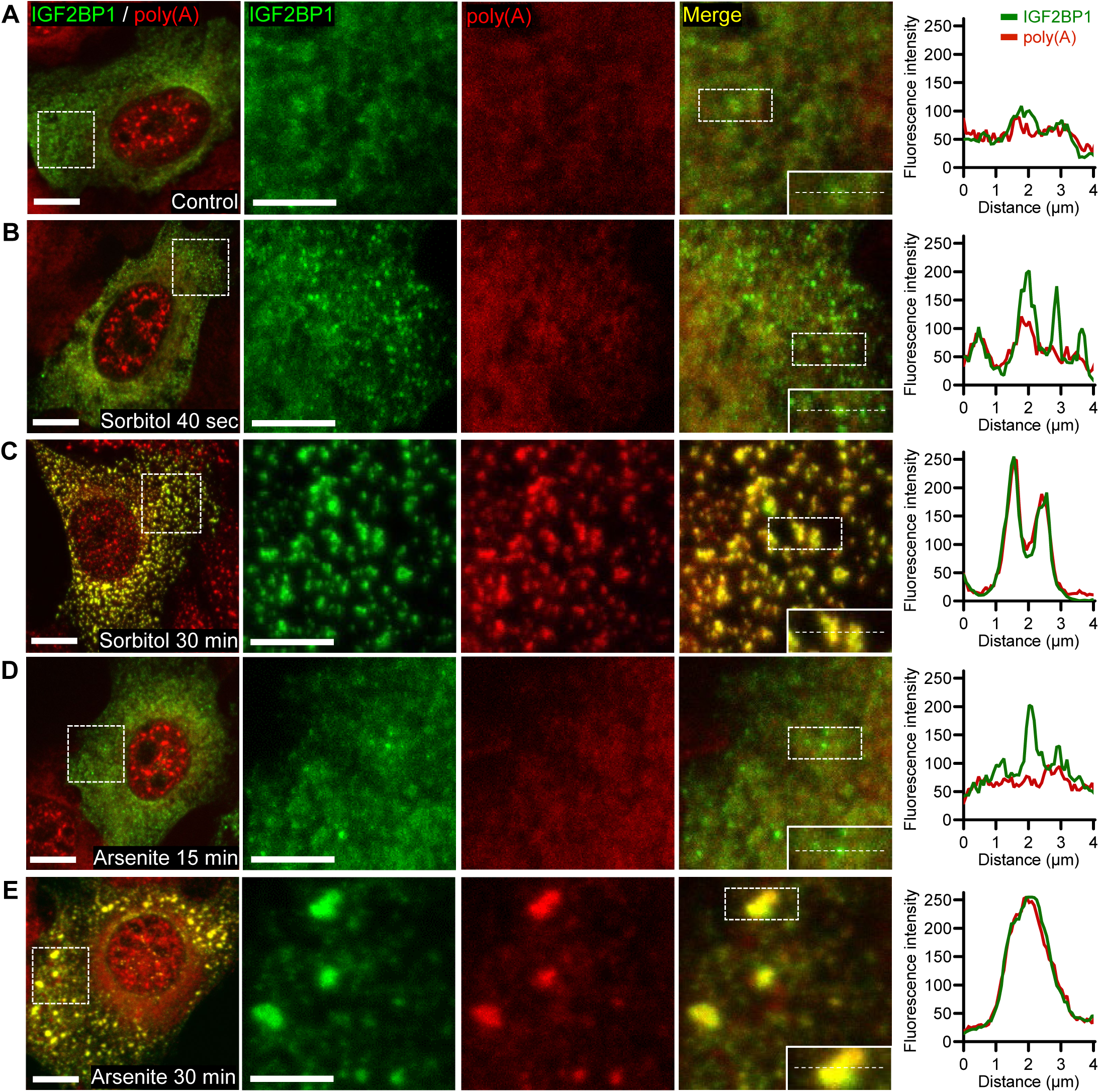
Poly(A) mRNA was rapidly associated with IGF2BP1 clusters and colocalized with SGs during osmotic stress. **(A-E)** Dual-color confocal microscopy images of GFP-IGF2BP1 (green) and poly(A) mRNA (red) in an U2OS stable cell line. Cells were either untreated (A) or treated with 375mM sorbitol (B-C) or 0.5mM sodium arsenite (D-E) for the indicated periods before fixation with 4% paraformaldehyde. poly(A) mRNA was detected by FISH using a Cy3-labeled oligo(dT) probe. The left column presents the whole cell images. The next three columns to the right display the enlarged images of the areas marked by the dashed boxes within the corresponding whole-cell image in the same row. All scale bars are 10 μm for the left column and 5 μm for the enlarged images. Insets at the lower-right corner of each merged image are the enlarged images of the areas marked by the dashed boxes within the same image. The white dotted lines in the insets indicate the locations at which dual-color fluorescence intensities were measured. Plots of dual-color fluorescence intensities GFP-IGF2BP1 (green) and poly(A) mRNA (red) across individual clusters or SGs are shown in the rightmost column.

Next, we performed FISH using Cy3-labeled oligo-dT probes in the stable cell line to visualize poly(A) mRNA in conjunction with GFP-IGF2BP1 at different stress conditions by confocal laser scanning microscopy. We acquired images from different samples using identical imaging parameters (see Materials & Methods) and analyzed cells with comparable protein expression levels. Prior to osmotic stress, poly(A) mRNA displayed a speckle-like pattern in the nucleus and a generally diffuse pattern with some clusters in the cytoplasm (Figure 6A). We observed multiple weak clusters of GFP-IGF2BP1 in the cytoplasm in addition to a diffuse localization before stress (Figure 6A), which supported previous findings that IGF2BP1/ZBP1 existed as granules in the cytoplasm (Farina et al., 2003; Jonson et al., 2007; Nielsen et al., 2002) and suggested that preexisting IGF2BP1 clusters in the cytoplasm might rapidly form core structures to initiate SG assembly in response to osmotic stress. At 40 sec after osmotic stress, GFP-IGF2BP1 formed numerous clusters that were larger in size and showed higher fluorescence intensities than those weak clusters prior to stress (Figure 6B), consistent with our live cell imaging results (Figure 1A). Interestingly, poly(A) mRNA also rapidly formed clusters upon osmotic stress, some of which were colocalized with IGF2BP1-marked clusters (Figure 6B). Clustering of poly(A) mRNA became increasingly evident at 3min, 5 min and 10 min after osmotic stress and those clusters were mostly colocalized with GFP-IGF2BP1 (Figure 6 – figure supplement 2). At 30 min after sorbitol treatment, strong poly(A) mRNA FISH signals were detected at SGs and were colocalized with GFP-IGF2BP1 (Figure 6C). Our findings thus indicated that a portion of poly(A) mRNA was rapidly recruited to IGF2BP1-marked clusters upon osmotic stress, and that the recruitment of poly(A) mRNA to SGs steadily increased during the assembly process.

The recruitment of poly(A) mRNA to SGs was markedly slower during oxidative stress. At 15 min after arsenite treatment, although many IGF2BP1-marked granules were formed in the cytoplasm, FISH signals of poly(A) mRNA at these granules were quite weak (Figure 6D). In contrast, at 30 min and 50 min after arsenite treatment, we detected large and dense poly(A) mRNA clusters that were colocalized with GFP-IGF2BP1 (Figure 6E, Figure 6 – figure supplement 2). Therefore, the recruitment of poly(A) mRNA to SGs during oxidative stress followed the clustering of SG proteins such as IGF2BP1, and also steadily increased during the assembly process.

### Analysis of SG formation by dual-color Stochastic Reconstruction Microscopy (STORM)

Previous studies indicated that mRNA molecules were present as distinct sub-clusters within SGs, and thus poly(A) mRNA was thought to be a component of SG core structures (Jain et al., 2016; Khong & Parker, 2018). Our study suggested that IGF2BP proteins might be components of SG cores because their clustering preceded the recruitment of other SG proteins during osmotic stress. In order to differentiate IGF2BP proteins and poly(A) mRNA as SG core components, we would need to analyze the spatial organizations of poly(A) mRNA and IGF2BPs before and during SG assembly. Hence, we generated a stable U2OS cell line expressing mEos2-IGF2BP1 and performed FISH to detect poly(A) mRNA using Alexa Fluor 647-tagged oligo-dT probes. We then imaged mEos2-IGF2BP1 and poly(A) mRNA FISH signals from U2OS cells at different stages of osmotic stress using dual-color STORM (Rust, Bates, & Zhuang, 2006) (Figure 7 – figure supplement 1). First, we detected multiple mEos2-IGF2BP1 clusters (100-200 nm in diameter) in the cytoplasm prior to stress (Figure 7A), similarly as GFP-IGF2BP1 clusters observed by confocal microscopy (Figure 6A). In contrast, poly(A) mRNA was mostly dispersed in the cytoplasm and was mostly not associated with mEos2-IGF2BP1 clusters prior to stress (Figure 7A), suggesting that the majority of poly(A) mRNA was not bound to IGF2BP1 before stress. Next, we found that at 40 sec after sorbitol treatment, IGF2BP1 coalesced into larger clusters (∼200-400 nm in diameter, some clusters were elongated, see Figure 7B), while some poly(A) mRNA were located at the periphery of IGF2BP1-marked clusters (Figure 7B). This finding was consistent with the confocal microscopy result that poly(A) mRNA was colocalized with some GFP-IGF2BP1 clusters upon osmotic stress (Figure 6B). At 30 min after sorbitol treatment, large (over 500 nm in diameter) and dense IGF2BP1 granules were formed (Figure 7C). Interestingly, poly(A) mRNA was detected as multiple dense clusters (∼100 nm in diameter) associated with both the interior and the periphery of each IGF2BP1 granule (Figure 7C). We note that because IGF2BP1 formed dense granules after extended stress, mEos2 fluorescence signals were concentrated at these granules and it became difficult to detect blinking fluorescence signals at the granules. Thus, we only analyzed those granules with detectable mEos2-IGF2BP1 blinking signals (see Video 18 for the acquired raw data of dual-color STORM imaging).

**Figure 7.**
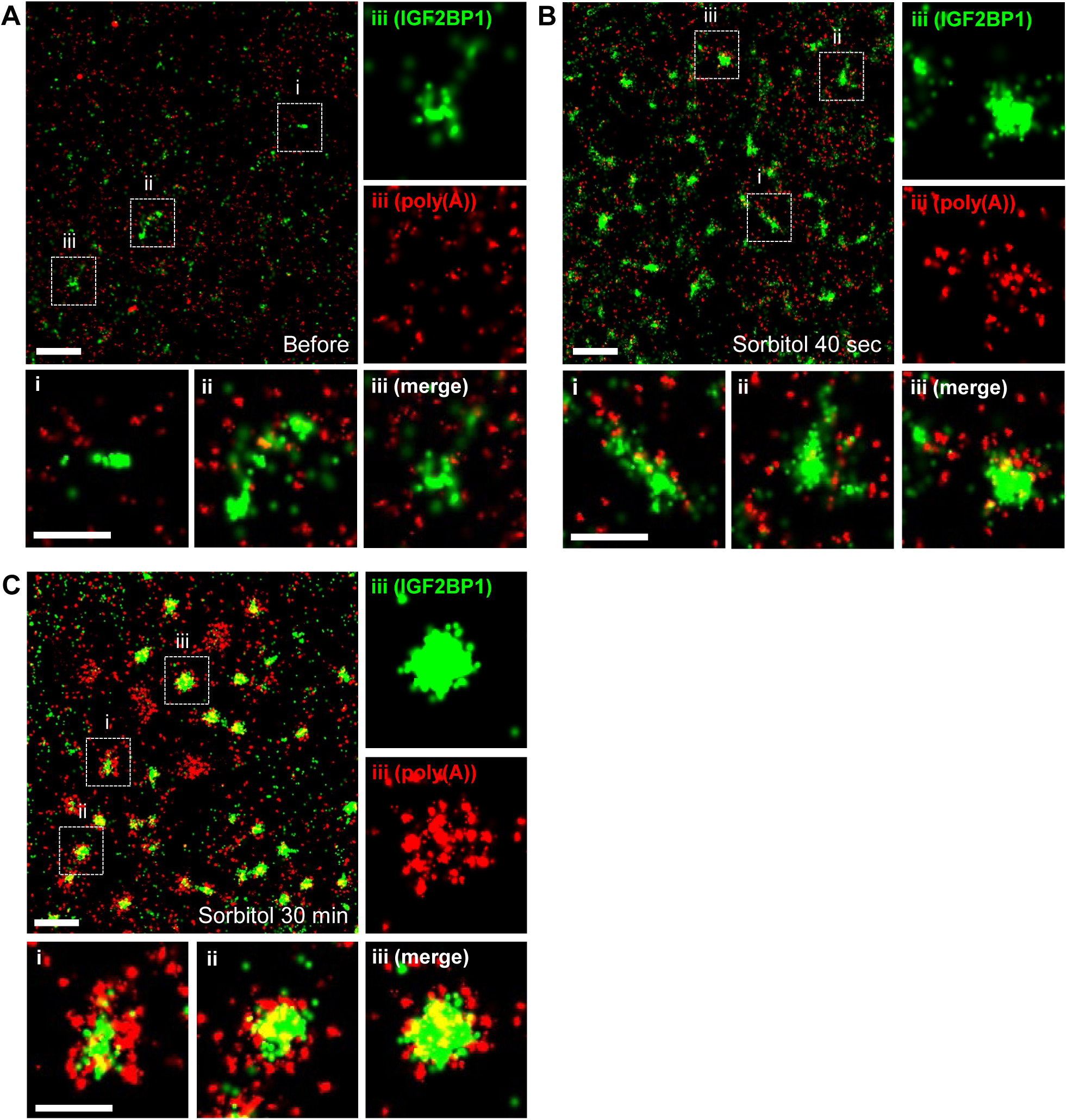
Visualizing the recruitment of IGF2BP1 and poly(A) mRNA to SGs during osmotic stress by dual-color STORM. **(A-C)** Dual-color STORM images of mEos2-IGF2BP1 (green) and poly(A) mRNA (red) in a U2OS stable cell line. Cells were either untreated (A) or treated with 375 mM sorbitol for 40 sec (B) or 30 min (C). Cells were then fixed with 4% paraformaldehyde and poly(A) mRNA were detected by fluorescence in situ hybridization (FISH) using an Alexa Fluor 647-labeled oligo(dT) probe. Images were acquired and analyzed at a Nikon N-STORM super-resolution microscope system. Within each panel, three dashed boxes were marked on the upper-left image (shown as i, ii, iii), each of which contains a representative IGF2BP1-marked cluster or SG and is enlarged in the corresponding lower row images. Single-color STORM images of mEos2-IGF2BP1 (green) and poly(A) mRNA (red) in box iii are shown above the merged image. Scale bars are 1 μm for the upper-left image and 500 nm for the enlarged images in the lower row of each panel.

**Figure 8.**
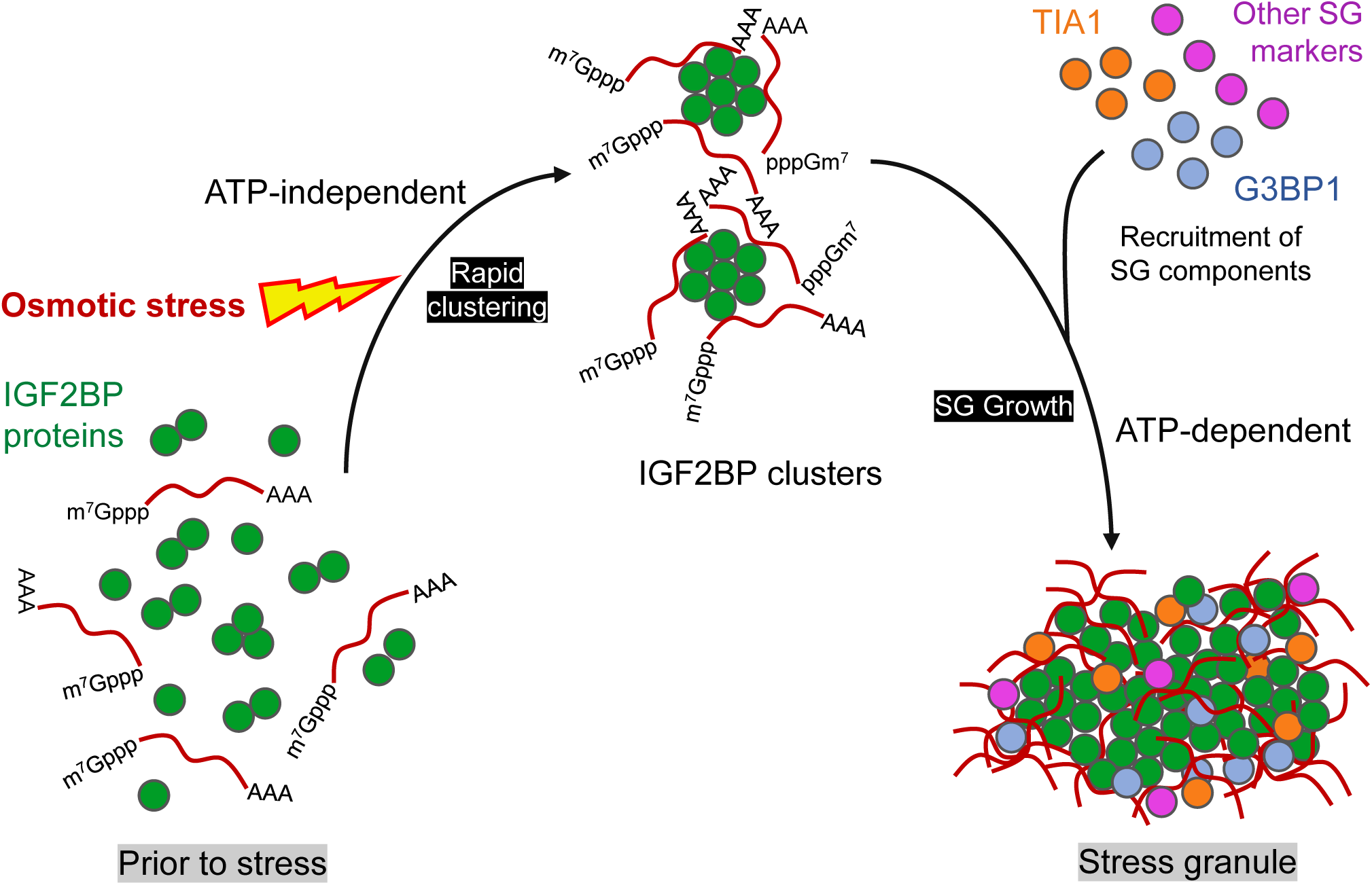
A cartoon model of SG assembly during osmotic stress in human cells.

In summary, our dual-color STORM imaging has confirmed the following findings by confocal microscopy: 1) IGF2BP1 exists as numerous small clusters in the cytoplasm prior to stress, and these small clusters are mostly unassociated with poly(A) mRNA; 2) A portion of poly(A) mRNA becomes associated with IGF2BP1-marked clusters at early stage of osmotic stress (i.e., 40 sec). Moreover, STORM imaging has revealed that poly(A) mRNAs form multiple clusters within a single SG at the late stage (i.e., 30 min) of osmotic stress. These clusters of poly(A) mRNA are associated with IGF2BP1-marked granules both at the periphery and in the interior. Therefore, SGs formed during osmotic stress are highly heterogeneous organelles, in which poly(A) mRNA and IGF2BP1 are organized as distinct sub-structures.

## Discussions

Our study has used live cell imaging to reveal a novel SG assembly process during osmotic stress. Here we unexpectedly found that IGF2BP proteins rapidly formed multiple clusters in the cytoplasm upon osmotic stress, followed by the recruitment of other SG proteins such as G3BP1 and TIA1. In contrast, all three proteins were recruited to SGs following a similar kinetics during oxidative stress. Therefore, SG assembly during osmotic stress proceeds in a distinct pathway from SG assembly during oxidative stress. Interestingly, we found that the clustering of IGF2BP proteins upon osmotic stress was ATP-independent, while the recruitment of G3BP1 and TIA1 to SGs was ATP-dependent. Some ATP-dependent processes during SG assembly may include mRNA remodeling and protein chaperone activities. Understanding these energy-consuming processes and the roles of relevant proteins will provide more insights into SG biogenesis.

Imaging studies have revealed the heterogeneous organization of an SG, in which a single SG contains multiple core structures marked by high concentrations of mRNAs or proteins (Jain et al., 2016; Wheeler et al., 2016). How core structures are formed during SG assembly has not been well understood. Our observations suggested that IGF2BP clusters might serve as “precursor granules” to subsequently recruit mRNA and other SG proteins and to fuse into larger SGs. In line with this hypothesis, dual-color STORM imaging revealed that SGs formed after extended osmotic stress contained a single dense granule of IGF2BP1 and multiple poly(A) mRNA clusters located within or at its periphery (Figure 7C). Given that IGF2BP proteins function to enhance the stability of several oncogenic mRNAs (Huang et al., 2018) and that IGF2BP expression often correlates with poor prognosis in cancer (Bell et al., 2013; Degrauwe, Suva, Janiszewska, Riggi, & Stamenkovic, 2016), it would be interesting to examine whether the rapid clustering of IGF2BP proteins occurs in certain malignant cell types under stress and whether this process might be targeted to downregulate oncogenic mRNA expression.

A conundrum potentially related to our findings is the different residence times of SG proteins observed in living cells. While several SG proteins such as G3BP1 and TIA1 remain associated with SG for only a few seconds, some other SG proteins including IGF2BP1 reside at SGs on the time scale of a minute or longer (Bley et al., 2015). Stable associations of SG proteins suggested that diffusion of these proteins out of SGs was delayed as a result of homotypic or heterotypic interactions with SG components. Analyzing these interactions occurring within SGs as diffusion barriers will help to understand the heterogeneous organization of SGs. Therefore, it would be interesting to reveal the specific domains of IGF2BP proteins responsible for mediating stable association with SGs. Furthermore, whether other stably associated SG proteins may be components of “precursor granules” formed upon osmotic stress remains to be examined.

Our STORM experiments revealed that IGF2BP1 proteins formed clusters on the order of ∼100nm in size prior to stress and these clusters rapidly coalesced into microscopically visible clusters (200-400 nm) upon osmotic stress. These findings are consistent with recent results that some SG proteins were detected as small clusters in the cytoplasm prior to stress (Narayanan et al., 2019). Preexisting protein clusters indicate highly cooperative associations between monomers and may serve as effective sensors to hyperosmolarity in the cell or other environmental signals. Characterizations of these preexisting protein clusters such as their molecular identities, spatial distributions relative to known cellular structures, diffusion and/or transport dynamics are needed. Development and application of high-resolution, high-speed live cell imaging methods will be essential for such studies.

## Materials and Methods

### Cell Culture and Transfection

U2OS cell line was obtained from Chinese Academy of Sciences, Kunming Cell Bank (http://www.kmcellbank.com). U2OS cells were cultured in low glucose Dulbecco’s Modified Eagle’s Medium (DMEM, ThermoFisher), 10% fetal bovine serum (FBS, Hyclone), 1% penicillin/streptomycin (ThermoFisher) and 1% GlutaMax (Gibco) in a CO_2_ incubator at 37°C. Cells were transfected with Lipofectamine 3000 (ThermoFisher) according to the manufacturer’s instructions. Transfection was performed in growth media without antibiotics. Cells were transfected for 8 hours and were subsequently cultured in normal growth media for 12 hours before analyses.

### cDNA Cloning and Plasmid Construction

Human IGF2BP1, IGF2BP2, IGF2BP3, G3BP1 and TIA1 cDNA were amplified by RT-PCR and cloned into pAcGFP-C1, pEGFP-N1, pmCherry-C1 and pmEos2-C1 (generated in a previous study (Yao, Fetter, Hu, Betzig, & Tjian, 2011)). The open reading frames (ORFs) of IGF2BP1, IGF2BP2, IGF2BP3, G3BP1 and TIA1 coding region sequence amplified by RT-PCR was chosen from the following NCBI Reference Sequences: NM_006546.4, NM_006548.6, NM_006547.3, NM_005754.3 and NM_022037.4, respectively.

U2OS total RNA was extracted with a ReliaPrep RNA cell Miniprep System (Z6011, Promega) according to the manufacturer’s instructions and was used as the template for RT-PCR using Superscript IV One-Step RT-PCR System (Invitrogen, 12594-025). Briefly, 1 ng RNA template was mixed with the primer pairs (described in Table S1) and the enzyme mix at a final volume of 50 µl. RT-PCR reaction was then performed according to the manufacturer’s instructions. For cloning IGF2BP1 truncation mutants, the indicated primer pairs were mixed with 1ng of pAcGFP-C1-IGF2BP1 plasmid in 1X Phusion High-Fidelity DNA Polymerase (F530L, ThermoFisher) mix at a final volume of 50 µl. PCR reactions were performed according to the manufacturer’s instructions.

RT-PCR or PCR products were cleaned up with QIAquick PCR purification kit (28106, QIAGEN) or QIAquick Gel Extraction kit (28706, QIAGEN). Purified PCR products and fluorescent protein expression plasmids were digested with the corresponding restricted enzymes (ThermoFisher, see Table S1 for details) and then cleaned up with a PCR purification kit. The digested fluorescence vectors were then treated with calf intestine phosphatase (CIP, M0290, NEB). The digested PCR products and CIP-treated vectors were ligated with T4 DNA ligase (M0202, NEB) at 16°C on a PCR machine overnight. All cloned ORFs and in-frame fusion with fluorescent protein coding sequences were confirmed by sequencing.

### Stable Cell Line Generation

U2OS cells were transfected with pAcGFP-C1-IGF2BP1 plasmid or pmEos2-C1-IGF2BP1 plasmid. After 48 hours of transfection, the culture media was replaced with selection media containing low glucose DMEM, 10% FBS, 1% GlutaMax and 400 μg/ml G418 sulfate (Yeasen Biotechnology, Shanghai, China). Cells were subsequently cultured in selection media for 14 days with media change every three days. Treatment of mock transfected U2OS cells with selection media was performed to ensure the efficacy of selection. G418-resistant cells after 14-day selection were collected and used for further experiments.

### Live Cell Imaging and Cell Treatment

Live cell imaging was performed at a Nikon Eclipse Ti2-E fluorescence microscope within a stagetop incubator (Tokai Hit model STX) for controlling temperature, humidity and the CO_2_ level during image acquisition. U2OS cells were seeded on a glass bottom dish (D35-20-1-N, Cellvis, Mountain View, CA, USA) and were equilibrated at 37°C and 5% CO_2_ in the humidified stagetop incubator for 15 min before image acquisition. A 60X oil-immersion objective (CFI Plan Apochromat Lambda, NA 1.40, Nikon) was used for live cell imaging. A 1.5X magnifier lens was placed in the light path during live cell imaging. Images were acquired with a DS-Qi2 CMOS camera (Nikon) using NIS-Element software (v5.1, Nikon). A 32X neutral density filter was applied after the fluorescence mercury lamp (C-HGFI, Nikon) to attenuate the excitation light. GFP fluorescence was collected through a C-FL-C FITC filter cube (MBE44725, Nikon) at 50ms exposure at 20X analog gain. mCherry/RFP fluorescence was collected through a C-FL-C Texas Red filter cube (MBE46105, Nikon) at 100ms exposure at 20X analog gain. Single-color or dual-color images of the same cell were acquired under the parameters described above at 10-second intervals for up to 50 minutes after stress induction. The TI2-N-ND-P perfect focus unit (Nikon) was used to maintain image focus during acquisition. After acquiring images for one to five minutes, the culture media was replaced with growth media containing 375mM sorbitol (A610491, BBI Life Sciences, Shanghai, China) for osmotic stress or with growth media containing 0.5mM sodium arsenite (S7400, Sigma) for oxidative stress. For ATP depletion, cells were treated with media containing 375mM sorbitol, 200mM 2-DG (D8375, Sigma) and 100µM CCCP (C2759, Sigma). A custom-made injection system was used to ensure rapid withdrawal of the old culture media and application of the stress-inducing media.

### Cell Fixation and Widefield Fluorescence Microscopy

U2OS cells were seeded on coverslips (12-541-B, Fisherbrand) placed in a 6-well plate and were transfected with plasmids prior to treatment and fixation (images are presented in Figure 5). After treatment with stressors, coverslips were washed once with 1X PBS and cells were fixed with 4% paraformaldehyde (28908, ThermoFisher) in 1X PBS for 10 min. Cells were then washed three times with 1X PBS and mounted on a clean glass slide (PC3-301, MeVid, Jiangsu, China) with Fluoromount-G mounting media (0100-01, Southern Biotech, Birmingham, AL, USA). U2OS cells stably expressing GFP-IGF2BP1 were washed twice with 1X PBS, stained with Hoechst 33342 (1:10,000 in 1X PBS, ThermoFisher) for 10 min, washed again three times with 1X PBS and mounted on a clean glass slide with Fluoromount-G mounting media (images are shown in Figure 6 – figure supplement 1B). Images of fixed cells were acquired at a Nikon Ti2-E widefield fluorescence microscope using a 60X oil-immersion objective and a DS-Qi2 CMOS camera.

### FISH and Confocal Microscopy

U2OS cells stably expressing AcGFP-IGF2BP1 were seeded on coverslips for FISH and confocal laser scanning microscopy. Poly(A) mRNA was detected using the following FISH protocol (https://openwetware.org/wiki/Poly_A_RNA_in_situ_protocol). Briefly, after treatment with stressors, U2OS cells were washed once with 1X PBS and fixed with 4% paraformaldehyde in 1X PBS for 10 min. Cells were then permeabilized with 100% methanol prechilled at -20°C for 10 min and rinsed with 1M Tris (pH 8.0) for 5 min. Cells were hybridized with 70 ng Cy3-labeled 30-mer oligo(dT) probes (synthesized by Genscript, Nanjing, China) overnight at 37°C. After hybridization, cells were washed once with 4X SSC buffer and then once with 2X SSC buffer. Cells were then washed once with 1X PBS, stained with Hoechst 33342 (1:10,000 diluted in 1X PBS) for 10 min, and mounted on a clean glass slide with Fluoromount-G mounting media. Fluorescence images of Cy3-oligo(dT) and GFP-IGF2BP1 were acquired on a Nikon A1+ confocal microscope using a 100X oil-immersion objective lens (HP Apo TIRF 100xH, 1.49 NA, Nikon). Confocal images were acquired at 2.0 AU pinhole (111.1 μ m), 1024x1024 image size, a zoom factor of 1.6 (pixel size 80 nm). 13 image frames per z-stack with a 225 nm distance between frames were used to acquire fluorescence signals of a whole cell. Z-stacks images were processed into a max-intensity projection image using ImageJ.

### Image and Statistical Analysis

All images were processed and analyzed with ImageJ software (v1.52p). In Figure 1F, the mean fluorescence intensity was measured within SG areas of similar sizes at each time point during oxidative stress or osmotic stress. In Figure 6, dual-color fluorescence intensity was measured along the longer axis of a rectangular box of 54 x 4 pixels. The fluorescence intensity curves were generated by GraphPad Prism 8.2.1. Unpaired *t*-test analysis was performed using GraphPad Prism 8.2.1 to evaluate significant differences between two groups being compared, and *p* < 0.05 was considered statistically significant.

### Dual-color STORM Imaging

U2OS cells stably expressing mEos2-IGF2BP1 were seeded on a glass bottom dish (Cell-Vis) for FISH and dual-color STORM. oligo(dT) probes was labeled with Alexa Fluor 647 and then used to detect poly(A) mRNA by FISH. First, 20 µg of 30-mer oligo(dT) labeled with an NH_2_ group at its 5’-end (synthesized by Genscript, Nanjing, China) and 50 µg of Alexa Fluor 647 NHS ester (A37537, ThermoFisher) were mixed in the labeling buffer (0.1M sodium tetraborate (Sigma), pH 8.5) overnight at room temperature. After labeling, 10% volume of 3M sodium chloride and 2.5X volume of 100% cold ethanol was added to the labeling reaction. The mixture was then incubated at -20°C for 30 min and spun down at 12,000g for 30 min at 4°C. The supernatant was carefully removed and the pellet was rinsed twice with 70% cold ethanol. The pellet was air-dried and dissolved in 50 µl ddH_2_O.

FISH protocol was performed in the glass bottom dish. Cells were hybridized with 250 ng Alexa Fluor 647-labeled 30-mer oligo(dT) probes in 250 µl hybridization reaction. The STORM imaging buffer was prepared according to the protocol provided by Nikon. Glucose Oxidase (G2133, Sigma) and Catalase (C1345, Sigma) stock solution were dissolved in Buffer A (10 mM Tris, 50 mM sodium chloride, pH 8.0) with a final concentration of 70 mg/ml and 17 mg/ml, respectively. GLOX solution was prepared freshly by mixing Glucose Oxidase and Catalase stock solution at 4:1 ratio. The imaging buffer was freshly prepared by mixing 7 µl GLOX, 7 µl 2-mercaptoethanol (07604, Sigma) and 690 µl Buffer B (50 mM Tris, 10 mM sodium chloride, 10% Glucose, pH 8.0), and added into the glass bottom dish before STORM imaging. Fluorescence signals of mEos2-IGF2BP1 and Alexa Fluor 647-oligo(dT) were detected sequentially on a Nikon N-STORM microscope system using a 100X oil objective lens (APO TIRF, 1.49NA, Nikon) and an ORCA-Flash 4.0 V2 sCMOS camera (Hamamatsu) at 40ms exposure in the no-binning mode. mEos2-IGF2BP1 was excited with a 561nm laser and fluorescence was detected in the RFP channel. Photoconversion of mEos2 was triggered by a 405nm laser and the intensity of the 405nm laser was gradually increased until all mEos2 molecules were photoconverted and detected. For imaging poly(A) mRNA, Alexa Fluor 647-oligo(dT) signals were excited with a 640nm laser and fluorescence was detected in the Cy5 channel. “On”-“off” photoswitching of Alexa Fluor 647 occurred spontaneously and a 405nm laser was occasionally used to accelerate the photoswitching. Single molecule localization and image reconstruction were performed by Nikon NIS Element software (v5.1) with N-STORM plugin.

### Western Blot

Approximately 10^7^ cells from the U2OS stable cell line expressing GFP-IGF2BP1 were washed once with 1X PBS and lysed in RIPA buffer prepared according to (http://cshprotocols.cshlp.org/content/2017/12/pdb.rec101428.short). The cell lysate was collected and heated with loading buffer for 5 min at 95 °C. Proteins were resolved by SDS-PAGE on a 10% polyacrylamide gel and transferred to 0.2 µm nitrocellulose membrane on a semi-dry transblot (Trans-Blot Turbo, Bio-Rad). The membrane was blocked with 5% BSA (Sigma) in TBST (0.05% Tween20) and then probed with a rabbit anti-IGF2BP1 polyclonal antibody (Abcam 82968, raised against 36-85 amino acids of human IGF2BP1/IMP1) at 1:1000 dilution or a mouse anti-β-actin monoclonal antibody (sc-47778, Santa Cruz) at 1:1000 dilution overnight at 4°C. The membrane was then incubated with a goat-anti-rabbit secondary antibody conjugated to horseradish peroxidase (Life Technologies, 1:5000) or a goat-anti-mouse secondary antibody conjugated to horseradish peroxidase (Cell Signaling, 1:5000) at room temperature for 1 hour. Specific protein bands on the membrane were detected by the ECL detection reagent (34577, ThermoFisher) and visualized on a Tanon-5200 Chemiluminescent Imaging System (Tanon Science and Technology, Shanghai, China).

## Supporting information

Video 1. Rapid clustering of GFP-IGF2BP1 during osmotic stress.

Video 2. Rapid clustering of GFP-IGF2BP1 after adding hypertonic culture media.

Video 3. Recruitment of GFP-G3BP1 to SGs during osmotic stress.

Video 4. Recruitment of TIA1-GFP to SGs during osmotic stress.

Video 5. Recruitment of GFP-IGF2BP1 to SGs during oxidative stress.

Video 6. Recruitment of GFP-G3BP1 to SGs during oxidative stress.

Video 7. Recruitment of TIA1-GFP to SGs during oxidative stress.

Video 8. Rapid clustering of GFP-IGF2BP1 during osmotic stress and ATP depletion.

Video 9. Live cell imaging of GFP-G3BP1 during osmotic stress and ATP depletion.

Video 10. Live cell imaging of TIA1-GFP during osmotic stress and ATP depletion.

Video 11. Live cell imaging of GFP-IGF2BP1(RRM1-KH2) fragment during osmotic stress.

Video 12. Live cell imaging of GFP-IGF2BP1(RRM1-KH3) fragment during osmotic stress.

Video 13. Live cell imaging of GFP-IGF2BP1(KH1-4) fragment during osmoticstress.

Video 14. Live cell imaging of GFP-IGF2BP1(KH3-4) fragment during osmotic stress.

Video 15. Live cell imaging of GFP-IGF2BP1(KH1-2) fragment during osmotic stress.

Video 16. Live cell imaging of GFP-IGF2BP1(aa157-KH2) fragment during osmotic stress.

Video 17. Live cell imaging of GFP-IGF2BP1(RRM1-2) fragment during osmotic stress.

Video 18. Videos of detecting fluorescence intermittency of mEos2-IGF2BP1 and Alexa Fluor 647-(dT)30 in U2OS cells after treatment with 375mM sorbitol

## Acknowledgements

We thank Prof. Deyin Guo and Prof. Yuan Chen (Sun Yat-sen University) for critical reading of this manuscript. We would like to thank Yuanjun Guan (Sun Yat-sen University) and Nikon Instruments (Shanghai) Guangzhou Branch for help with 3D STORM experiments. This work was supported by Hundred Talent Plan of Sun Yat-sen University.

## Author Contribution

Wei-jie Zeng, Conceptualization, Methodology, Validation, Investigation, Formal Analysis, Visualization, Writing – original draft, Writing – review and editing; Chuxin Lu, Methodology, Investigation; Yuanyuan Shi, Methodology; Xinxin Chen, Methodology; Jie Yao, Conceptualization, Validation, Formal Analysis, Visualization, Writing – original draft, Writing – review and editing, Supervision, Funding acquisition, Project administration.

## Competing Interest

The authors disclose no competing interest.

---

**Figure 1 – figure supplement 1.**
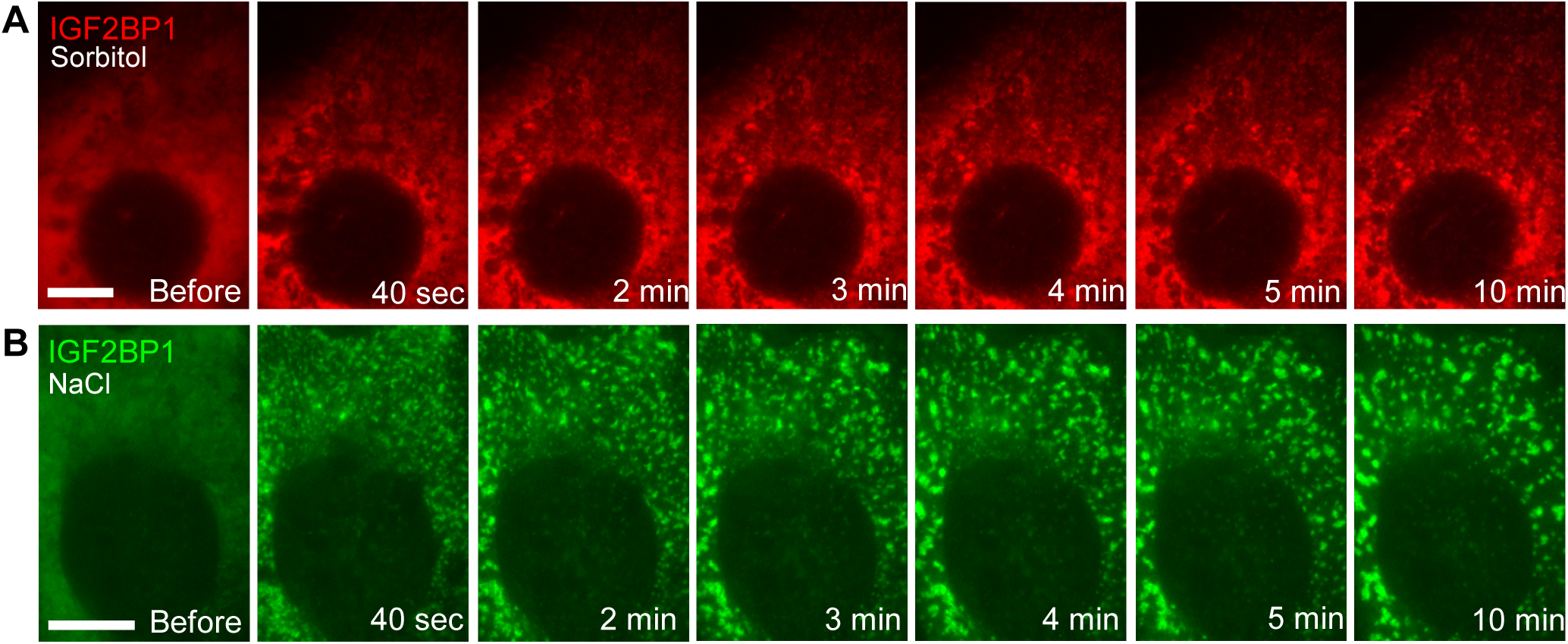
Rapid clustering of IGF2BP1 was observed from mCherry-fusion protein or after incubating cells with hypertonic growth media. **(A)** Time-lapse images of mCherry-IGF2BP1 in U2OS cells treated with 375mM sorbitol. **(B)** Time-lapse images of GFP-IGF2BP1 in U2OS cells treated with growth media supplemented with 100mM sodium chloride. Scale bars: 10 μm.

**Figure 1-figure supplement 2.**
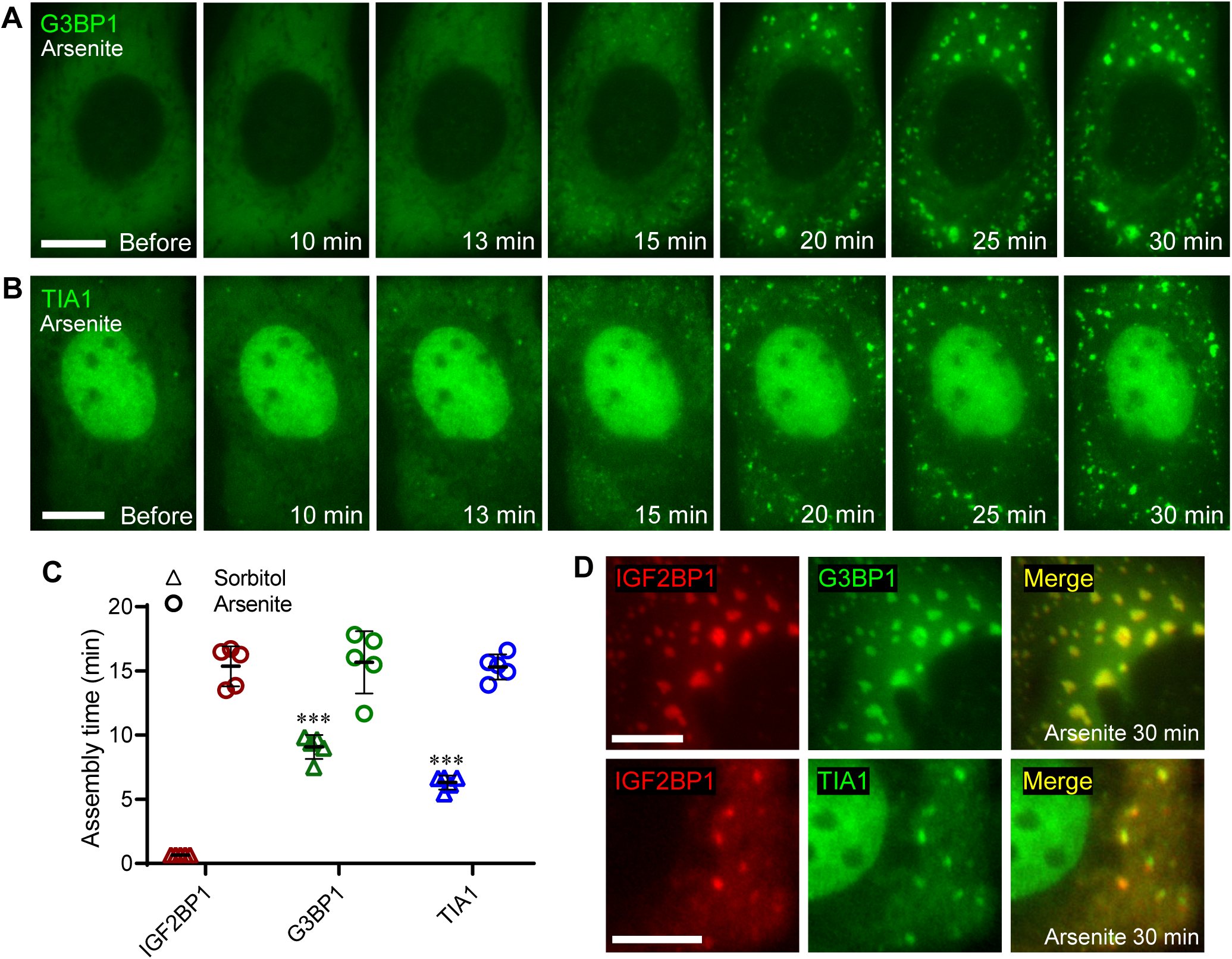
G3BP1 and TIA1 were recruited to stress granules concomitantly with IGF2BP1 during oxidative stress. **(A-B)** Time-lapse images of GFP-tagged G3BP1 (A) or TIA1 (B) in U2OS cells treated with 0.5mM sodium arsenite. Scale bars: 10 μm. **(C)** Plots of the time points of the first appearance of SGs marked by IGF2BP1, G3BP1 or TIA1 in U2OS cells after applying osmotic stress or oxidative stress. n = 5 in all groups. *** indicates *p* <0.001 in t-test between G3BP1 vs. IGF2BP1 or between TIA1 vs IGF2BP1 under sorbitol treatment. No significant difference was found between proteins under arsenite treatment. **(D)** Dual-color fluorescence images of U2OS cells co-expressing mCherry-IGF2BP1 and GFP-G3BP1 (upper) or TIA1-GFP (lower) after being treated with 0.5mM sodium arsenite for 30 min. Scale bars: 5 μm.

**Figure 4 – figure supplement 1.**
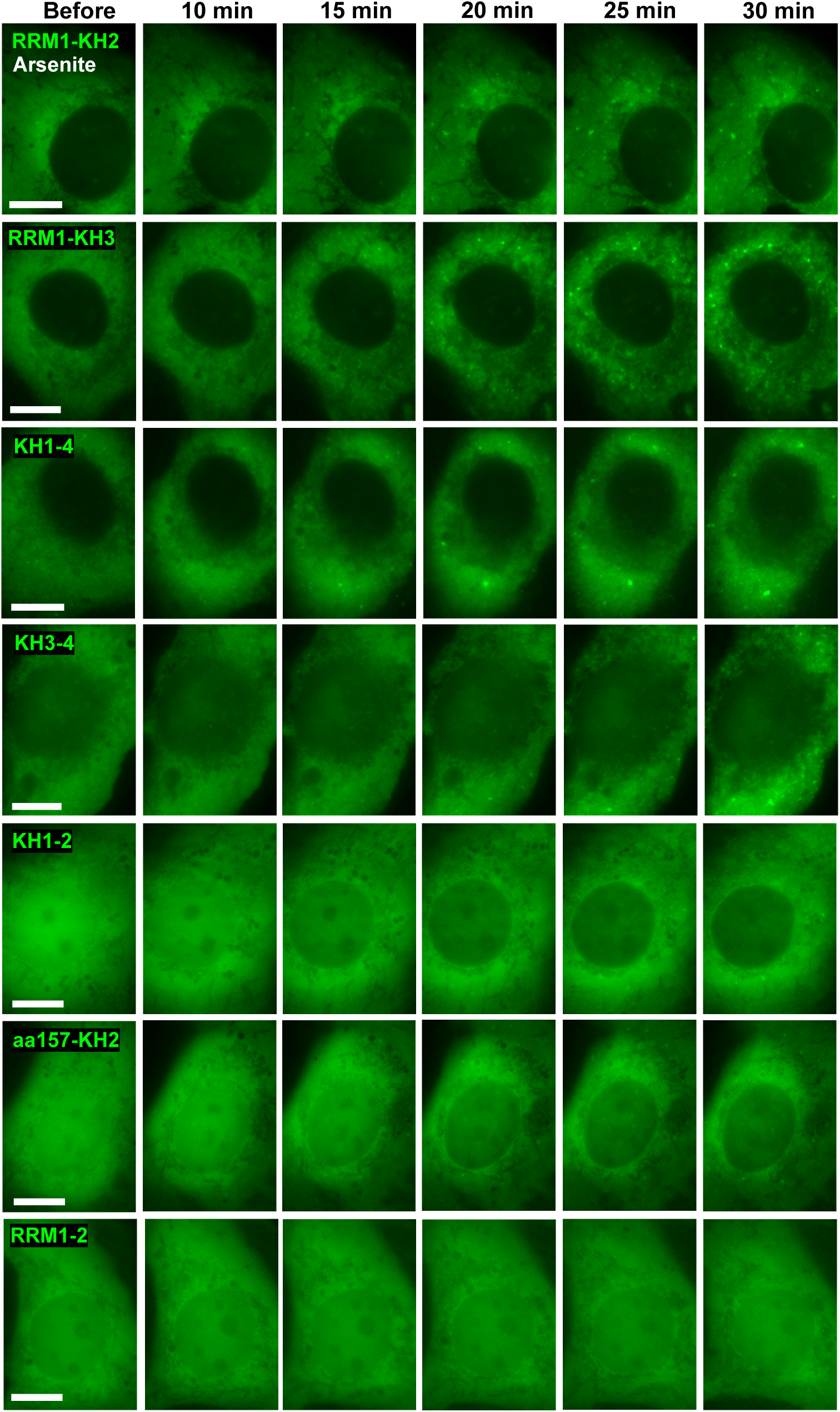
Formation of cytoplasmic aggregates by certain IGF2BP1 truncation mutants during oxidative stress. Each row shows time-lapse images of GFP-tagged IGF2BP1 truncation mutants (described in Figure 4A) in U2OS cells treated with 0.5mM sodium arsenite. Time intervals between applying oxidative stress and image acquisition are noted in each image. All scale bars are 10 μm.

**Figure 6 – figure supplement 1.**
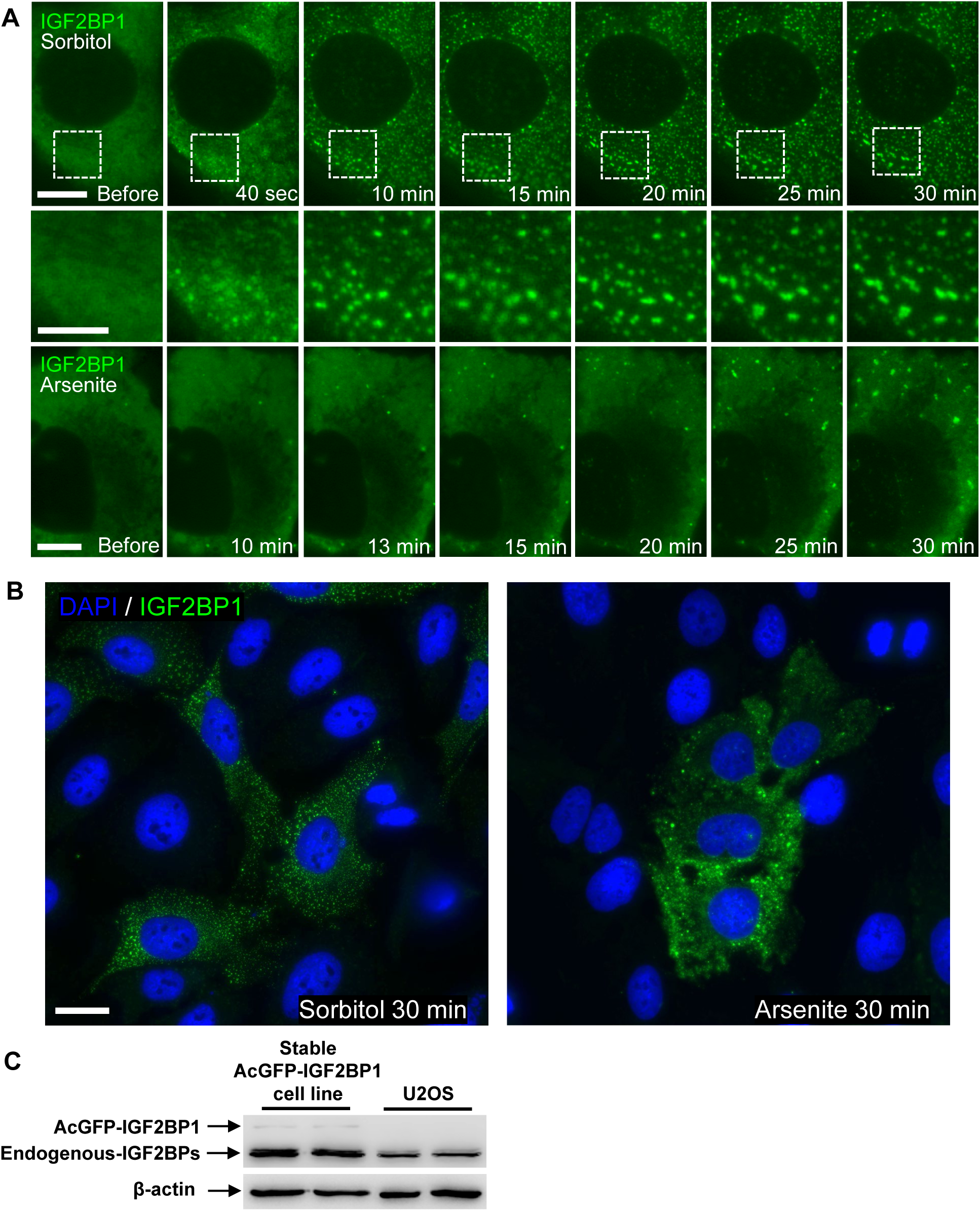
Rapid clustering and recruitment of IGF2BP1 into SGs during osmotic stress in a U2OS stable cell line. **(A)** Time-lapse images of a cell from the U2OS cell line stably expressing GFP-IGF2BP1 during osmotic stress (upper and middle rows) or oxidative stress (lower row). The middle row contains the enlarged images of the area marked by dashed boxes on each image above. Scale bars are 10 μm for upper and lower row images and 5 μm for the middle row images. Time points between applying stressor and image acquisition are noted on each image. **(B)** Fluorescence images of a U2OS stable cell line expressing GFP-IGF2BP1 (green) after treatment with 375mM sorbitol (left) or 0.5mM sodium arsenite (right). Nuclei were stained by Hoechst33342 (blue). Approximately 20-30% cells expressed GFP-IGF2BP1 at near-uniform levels. Scale bar: 20 μm. **(C)** Detecting the expression of GFP-IGF2BP1 and endogenous IGF2BPs in the U2OS stable cell line (shown in panel A, B) and endogenous IGF2BPs expression in wild type U2OS cells by western blot.

**Figure 6 – figure supplement 2.**
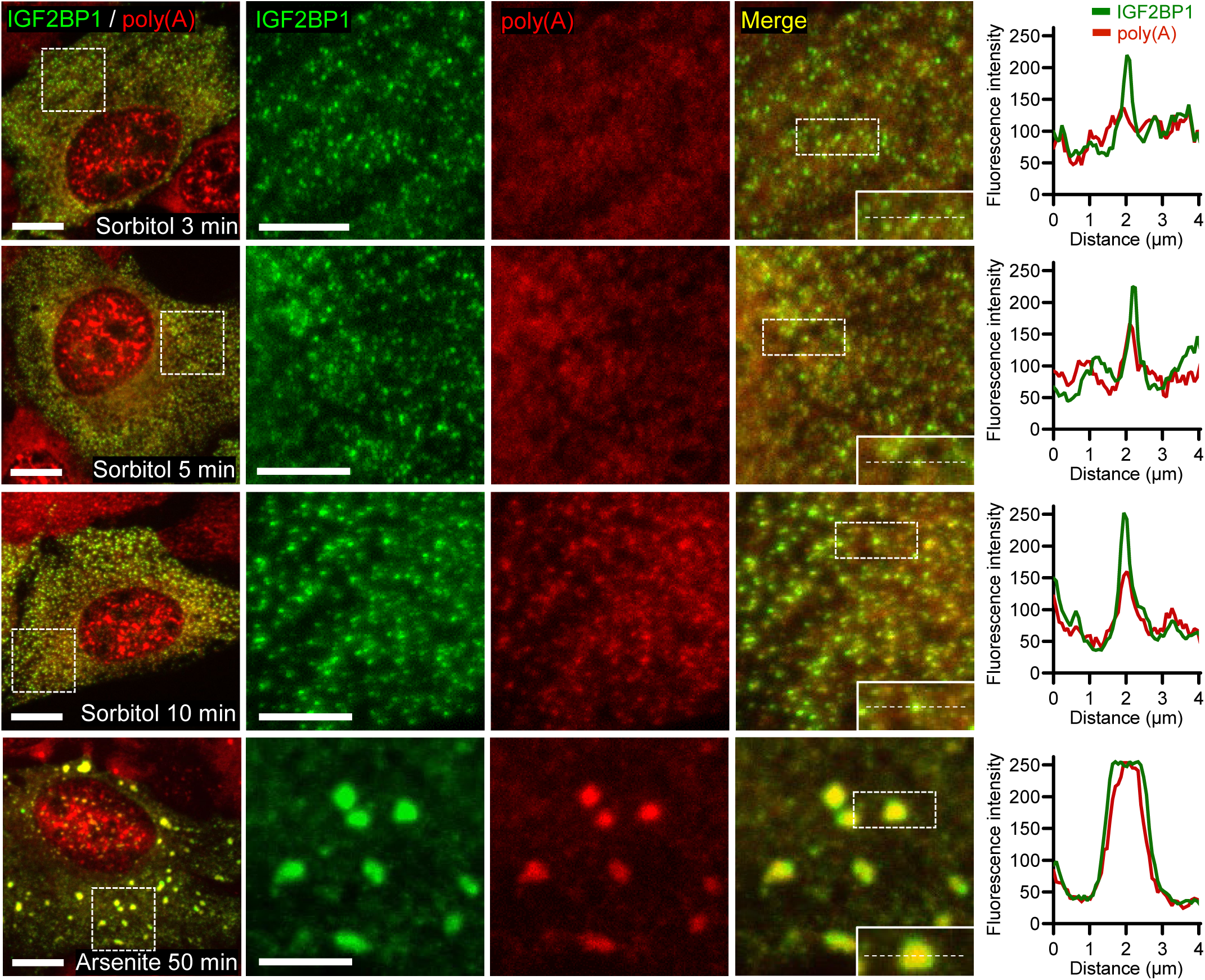
Dual-color confocal microscopy images of GFP-IGF2BP1 and poly(A) mRNA at additional time points during osmotic stress or oxidative stress. U2OS cells stably expressing GFP-IGF2BP1 (green) were treated with 375 mM sorbitol or 0.5 mM sodium arsenite for the time indicated on each image and were subsequently fixed and hybridized to Cy3-labeled oligo(dT)_30_ probes (red). Confocal images and fluorescence intensity data are presented as described in Figure 6. All scale bars are 10 μm for the left column and 5 μm for the enlarged images.

**Figure 7 – figure supplement 1.**
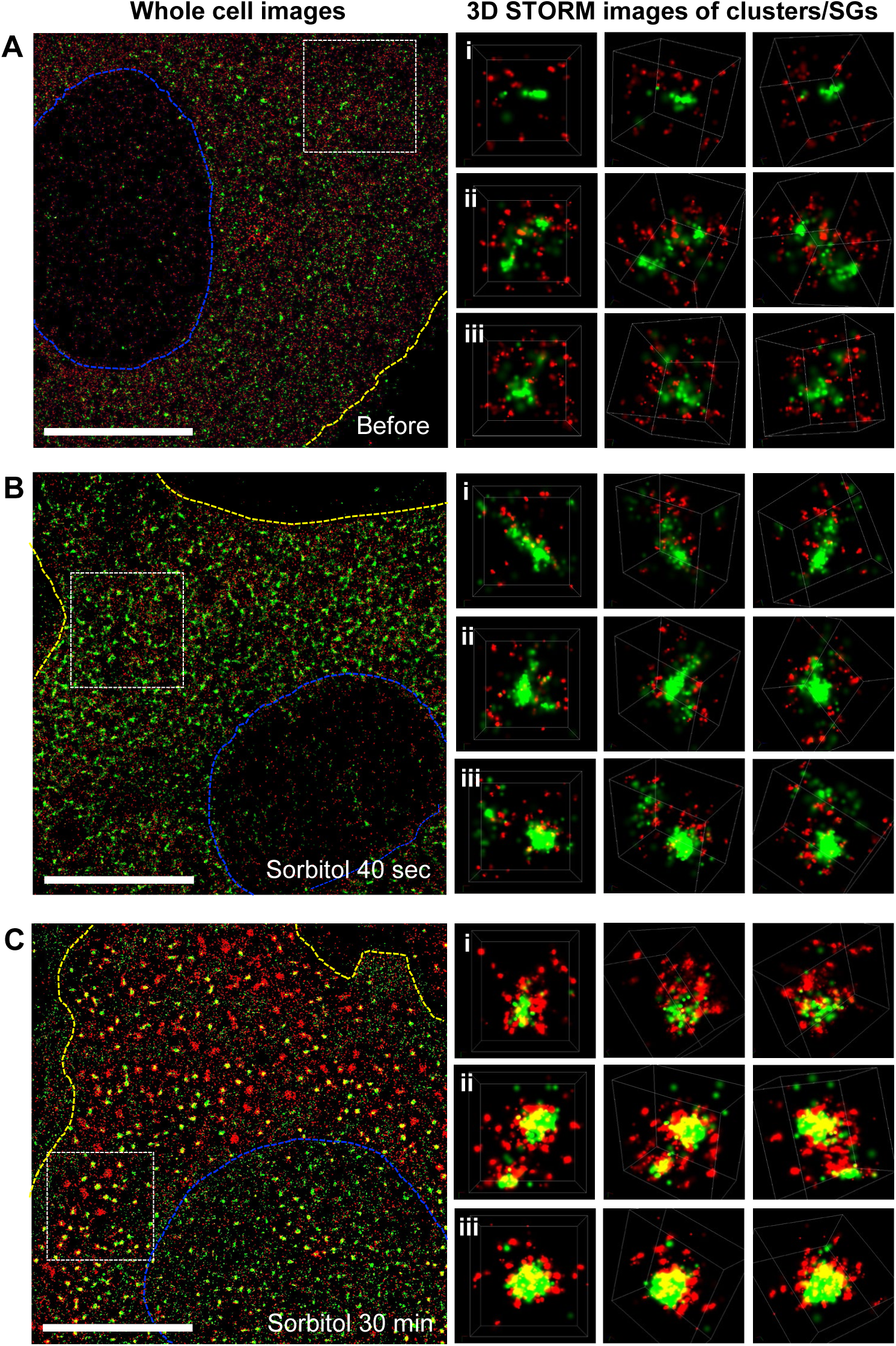
Dual-color 3D STORM images of IGF2BP1-containing clusters or SGs from cells shown in Figure 7. **(A-C)** The first column to the left presents the whole cell dual-color STORM images of mEos2-IGF2BP1 (green) and poly(A) mRNA (red) in a U2OS cell at the noted condition. Dashed blue lines indicate the nuclear borders and dashed yellow lines indicate the cell borders. The areas marked with dashed white box are shown as the upper-left images in Figure 7A-C, respectively. The three columns to the right are reconstructed dual-color 3D STORM images at three distinct angles of clusters/SGs from dashed boxes i, ii and iii shown in Figure 7A-C, respectively. Scale bars are 10 μm for the whole cell image.

**Table S1.**
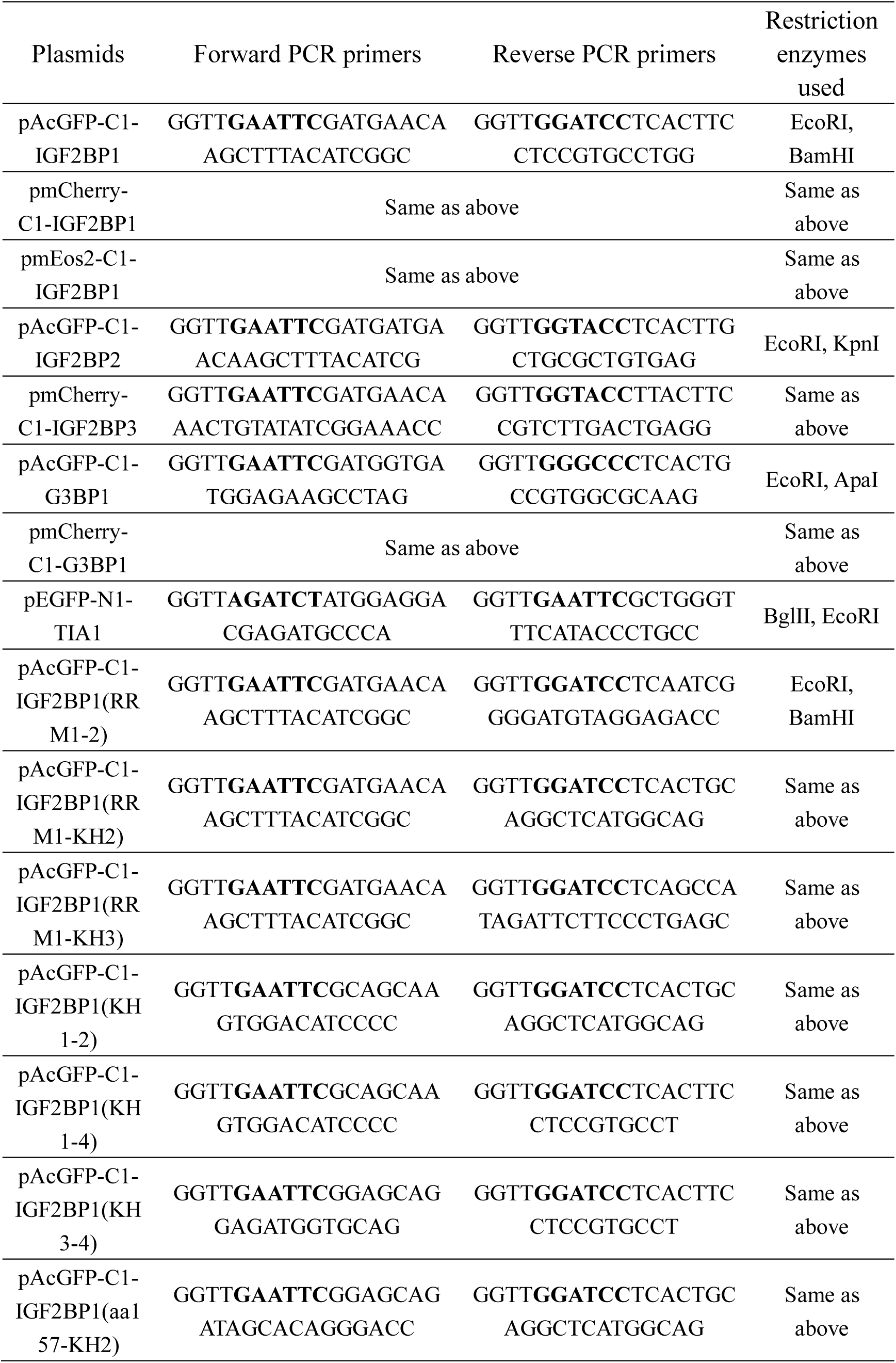
Plasmids used in this study

## Video Caption

**Video 1. Rapid clustering of GFP-IGF2BP1 during osmotic stress.** U2OS cells expressing GFP-IGF2BP1 were treated with 375mM sorbitol and imaged for 30 min. Images were acquired at 10 sec intervals. Related to Figure 1A.

**Video 2. Rapid clustering of GFP-IGF2BP1 after adding hypertonic culture media.** U2OS cells expressing GFP-IGF2BP1 were treated with growth media supplemented with 100mM sodium chloride and imaged for 30 min. Related to Figure 1 – figure supplement 1.

**Video 3. Recruitment of GFP-G3BP1 to SGs during osmotic stress.** Related to Figure 1B.

**Video 4. Recruitment of TIA1-GFP to SGs during osmotic stress.** Related to Figure 1C.

**Video 5. Recruitment of GFP-IGF2BP1 to SGs during oxidative stress.** U2OS cells expressing GFP-IGF2BP1 were treated with 0.5mM sodium arsenite and imaged for 30 min. Related to Figure 1D.

**Video 6. Recruitment of GFP-G3BP1 to SGs during oxidative stress.** Related to Figure 1 – figure supplement 2.

**Video 7. Recruitment of TIA1-GFP to SGs during oxidative stress.** Related to Figure 1 – figure supplement 2.

**Video 8. Rapid clustering of GFP-IGF2BP1 during osmotic stress and ATP depletion.** U2OS cells expressing GFP-IGF2BP1 were treated with growth media containing 375mM sorbitol, 200mM 2-deoxyglucose and 100µM carbonyl cyanide m-chlorophenyl hydrazone and imaged for 50 min. Related to Figure 2A.

**Video 9. Live cell imaging of GFP-G3BP1 during osmotic stress and ATP depletion.** Related to Figure 2B.

**Video 10. Live cell imaging of TIA1-GFP during osmotic stress and ATP depletion.** Related to Figure 2C.

**Video 11. Live cell imaging of GFP-IGF2BP1(RRM1-KH2) fragment during osmotic stress.** Related to Figure 4B.

**Video 12. Live cell imaging of GFP-IGF2BP1(RRM1-KH3) fragment during osmotic stress.** Related to Figure 4C.

**Video 13. Live cell imaging of GFP-IGF2BP1(KH1-4) fragment during osmotic**

**stress.** Related to Figure 4D.

**Video 14. Live cell imaging of GFP-IGF2BP1(KH3-4) fragment during osmotic stress.** Related to Figure 4E.

**Video 15. Live cell imaging of GFP-IGF2BP1(KH1-2) fragment during osmotic stress.** Related to Figure 4F.

**Video 16. Live cell imaging of GFP-IGF2BP1(aa157-KH2) fragment during osmotic stress.** Related to Figure 4G.

**Video 17. Live cell imaging of GFP-IGF2BP1(RRM1-2) fragment during osmotic stress.** Related to Figure 4H.

**Video 18. Videos of detecting fluorescence intermittency of mEos2-IGF2BP1 and Alexa Fluor 647-(dT)_30_ in U2OS cells after treatment with 375mM sorbitol for 30 min, acquired during dual-color STORM imaging.** Related to Figure 7.

## Notes

#### Summary of Updates

20191202 revised - complete version

